# Integrated Lipidomics and Nitro-Fatty Acid Profiling Link Adipose Redox Imbalance to Alzheimer’s Disease-Related Neurovascular Injury

**DOI:** 10.64898/2026.05.14.725257

**Authors:** Sukkum Ngullie Chang, Praveen Kumar Guttula, Kirti Agrawal, Elnaz Sheikh, William N. Beavers, Timothy D. Allerton, Manas Ranjan Gartia

## Abstract

Alzheimer’s disease (AD) is increasingly recognized as a systemic disorder in which peripheral metabolic and redox dysfunction affects neurovascular injury and amyloid pathology. However, the lipid–redox mechanisms linking adipose tissue dysfunction to AD are less explored. Here, we applied an integrated multi-omics and imaging approach to investigate adipose lipid remodelling that leads to nitric oxide (NO)–derived nitro-fatty acid modifications and their effects on β-amyloid and VEGF in the hippocampus of APP/PS1 mice. Targeted lipidomics revealed broad suppression of lysophospholipids and membrane phospholipids (LPC, PC, PE, PG) in gonadal white adipose tissue (gWAT), consistent with impaired membrane turnover and mitochondrial lipid deficiency. Untargeted lipidomics demonstrated accumulation of ceramides, triacylglycerols, monoacylglycerols, and phosphatidic acids, indicating lipotoxicity and disrupted lipid flux. Oxidized lipid mediator profiling showed increased 13-HODE, 12-HETE, and 14-HDHA. Further, Raman microscopy mapping revealed a shift from protective nitro-oleic acid toward increased nitration of polyunsaturated fatty acids. These lipid abnormalities coincided with increased adipose expression of redox and inflammatory markers, including NOX4 and TNF-α, and impaired mitochondrial redox metabolism assessed by fluorescence lifetime imaging (FLIM). Citrulline and nitrite treatments partially normalized adipose lipid–redox signatures. Citrulline restored phospholipid remodeling and nitro-oleic acid signaling, whereas nitrite preferentially enhanced stress-associated signaling lipids. Importantly, both interventions reduced hippocampal β-amyloid burden and restored VEGF expression, with citrulline producing the strongest neurovascular rescue. These findings identify adipose lipid–redox imbalance as a systemic contributor to neurovascular pathology in AD and highlight NO-directed metabolic modulation as a strategy to mitigate disease-associated lipid dysfunction.

## 1. Introduction

Alzheimer’s disease (AD) is widely recognized as a systemic disorder in which metabolic, redox, and inflammatory disturbances outside the brain contribute to neurodegeneration and vascular pathology [1–5]. Beyond classical amyloid-β (Aβ) and tau paradigms, peripheral tissues, particularly adipose depots, play an active role in modulating oxidative stress and lipid signaling that impact central nervous system (CNS) homeostasis [6, 7]. Obesity and high-fat diet (HFD) exposure are known epidemiological risk factors for AD. They create chronic low-grade inflammation, mitochondrial dysfunction, and altered lipid metabolism in adipose tissue [8, 9].

Emerging evidence implicates bidirectional metabolic communication between peripheral fat storage and the brain in the progression of neurodegenerative disease [10–12]. Adipocyte-derived cytokines, such as ceramides [13, 14], and lipid peroxidation products circulate systemically and can cross or influence the blood-brain barrier, promoting neuroinflammation and endothelial stress [12, 15, 16]. Mitochondrial dysfunction in adipose tissue amplifies this effect by impairing redox balance and fatty acid turnover, leading to the accumulation of lipotoxic species such as ceramides and oxidized phospholipids [17]. These metabolites are known to activate NADPH oxidase (NOX) and NF-κB–driven inflammatory cascades, reinforcing oxidative and nitrative stress both peripherally and centrally [18]. Despite these connections, the role of redox-driven lipid remodelling in adipose tissue in contributing to amyloid pathology remains poorly characterized.

Nitric oxide (NO) represents a critical connection in both metabolic and vascular regulation [19]. At physiological levels, NO and its derivatives (including nitro-fatty acids) preserve mitochondrial function, stimulate angiogenic signalling, and maintain endothelial homeostasis [20–22]. On the other hand, dysregulated NO signalling under oxidative conditions leads to aberrant nitration, loss of protective nitro-lipid species, and formation of pro-oxidant nitrated fatty acids [23]. Recent studies demonstrate that NO-directed dietary interventions such as nitrite and its precursor L-citrulline, can restore vascular NO bioavailability and mitigate metabolic inflammation [24, 25]. Yet, the extent to which such interventions normalize adipose lipid–redox pathways and influence neurovascular integrity in AD models exposed to metabolic stress remains largely unexplored.

The current understanding of NO biology in AD has primarily focused on endothelial dysfunction within the CNS. Early work established a link between reduced endothelial NO synthase activity and cerebrovascular compromise in Aβ pathology [26]. However, NO-mediated lipid signalling extends beyond the vasculature, influencing adipocyte differentiation, mitochondrial phospholipid remodelling, and interorgan lipid trafficking [27, 28]. The balance between protective nitro-oleic acid (nitro-OA) and harmful nitrated polyunsaturated fatty acids (nitro-CLA, nitro-AA, nitro-DHA) may therefore reflect the integrity of systemic redox–lipid regulation. Recent lipidomic advances have enabled simultaneous profiling of these nitro-lipid species, revealing that shifts toward pro-nitrative lipid signalling accompany both metabolic dysfunction and cognitive decline [29–31].

We performed comprehensive lipidomic, transcriptomic, and redox analyses to investigate lipid and lipid mediator changes in APP/PS1 and WT mice fed a high-fat diet (HFD). Further, Raman and fluorescence lifetime imaging microscopy (FLIM) experiments were performed to probe the effect of AD and HFD on NADH redox cycling and mitochondrial function. In addition, we investigated the effects of targeted NO-based therapeutics, such as citrulline and nitrite, on peripheral lipid-redox signalling and brain central neurovascular pathology. Finally, we demonstrated the effect of citrulline and nitrite on the hippocampal β-amyloid burden and VEGF expression. Our results could improve understanding of how adipose tissue metabolism drives neurovascular dysfunction in AD.

## 2. Materials and Methods

### 2.1. Animal model and experimental design

APPswe/PS1dE9 (APP) (n = 48 and wild-type (WT) littermate controls (n = 20) were purchased from Jackson Laboratory (Bar Harbor, ME) (age = 8 weeks, all female). All the animals were given a two-week acclimation period in a controlled environment to reduce stress and ensure consistent baseline health. Following acclimation (at age = 10 weeks), all received a high-fat diet (45% HFD; Research Diets 12451, New Brunswick, NJ) for 14 additional weeks (up to age = 24 weeks) to induce obesity and model metabolic dysfunction [32]. To evaluate the impact of nitric oxide-related supplementation, HFD-fed APP mice were further divided into three subgroups: Standard water (n = 18), Sodium nitrite (100 mg/L in water) (n = 18), and L-citrulline (400 mg/L in water) (n = 12). Mice had ad libitum access to their assigned diet and water. Food intake and body weight were measured weekly to track growth and dietary effects. Additionally, body composition (fat and lean mass) was assessed every two weeks using non-invasive nuclear magnetic resonance (NMR) imaging, providing detailed insight into physiological changes over time. At the end point, euthanasia was performed via isoflurane anesthesia followed by cardiac puncture. All procedures were approved by the Pennington Biomedical Research Center IACUC (approval # 20-098).

### 2.2. Glucose tolerance test

To further evaluate metabolic health, glucose tolerance testing (GTT) was performed in weeks 5, 15, and 25. This involved administering glucose and measuring blood glucose levels over time to assess how well the mice could regulate glucose levels. Following a 4-hour fasting period, glucose tolerance testing [32] was performed by first obtaining baseline blood samples via tail vein nick and measuring glucose levels using a handheld glucometer (Breeze2 glucometer, Leverkusen, Germany). Mice then received an intraperitoneal glucose injection at a dose of 3.5 g/kg lean body mass. Subsequent blood glucose measurements were collected from the tail vein at 15, 30, 45, 60, 90, and 120 minutes post-injection.

### 2.3. Hematoxylin and Eosin (H&E) Staining

Gonadal white adipose tissue (gWAT) samples were fixed in 4% paraformaldehyde and stored at 4°C for 24 hours. The tissues were embedded in paraffin and sectioned into blocks and allowed to solidify for 24 h, and sectioned at an 8 μm thickness using a microtome and attached to glass slides. Before the staining process, the slides were immersed in xylene for 30 min, followed by 100% ethanol (EtOH) for 3 min, 5% ethanol for 3 min, and 70% ethanol for 3 min. Next, the slides were washed with distilled water and stained with haematoxylin for 10 min, rinsed under running water and dipped in HCl for 30 s, rinsed under running water, dipped in ammonia for 30 s, and rinsed again under running water. Submerge the slides in eosin for 1 min and rinse under running water in a clean glass container. Repeat the ethanol series: 70% ethanol for 3 min, 95% ethanol for 3 min, and 100% ethanol for 3 min. Finally, place the slides in xylene, keeping them there unless they are immediately fixed with permount mounting medium and coverslips. The slides were examined under a light microscope at 4x, 20x, and 60x magnification.

### 2.4. Ex vivo lipolysis analysis

Gonadal white adipose tissue (gWAT) explants were rinsed with phosphate-buffered saline (PBS) and sectioned into individual pieces weighing approximately 20–30 mg. Multiple explants obtained from the same animal were utilized for lipolysis measurements. Tissues were incubated for 2 h at 37 °C in phenol red-free, low-glucose DMEM to assess basal lipolytic activity. Lipolysis was evaluated by quantifying glycerol released into the culture medium using a colorimetric assay kit (Sigma-Aldrich), with values normalized to the weight (g) of each tissue explant.

### 2.5. Lipidomics

#### Untargeted Lipidomics Experiment

The tissue preparation and experimental methodology were described previously [33]. In brief, the tissues were lysed in 250 μL of LC-MS grade methanol and homogenized with a Qsonica 500 until fully ground. Samples were kept cold throughout preparation. Protein concentration was assessed using the Bradford assay (Thermo Scientific) to determine tissue concentration, maintaining a 1:10 protein-to-tissue ratio. Based on the measured protein, volumes equivalent to 10 mg of tissue were transferred to new tubes. A 0.5 μg of EquiSPLASH LIPIDOMIX Quantitative Mass Spec Internal Standard (Avanti Polar, Birmingham, AL) was added, followed by vortexing for 1 min. After adding 200 μL of LC-MS grade methanol and vortexing for another minute, 400 μL of HPLC grade chloroform was added, vortexed, and kept on ice for 10 minutes, repeated three times. A 350 μL portion of the organic layer was transferred to a new tube and evaporated under a nitrogen stream. The samples were reconstituted in 50 μL of 40% methanol containing 0.1% formic acid. The Agilent 1260 Infinity II liquid chromatograph and Agilent 6230 Electrospray time-of-flight mass spectrometer were used for analysis. The capillary voltage was set at 4000 V, fragment voltage at 125 V, and nitrogen was used as the drying gas at 10 L/min at 325 °C. The mass range was 100–3000 m/z. A gradient program for chromatographic separation utilized an Agilent Poroshell 120 EC-C18 column at a flow rate of 400 μL/min. The mobile phase compositions were: A = 0.1% formic acid in water and methanol (60:40, v/v), and B = methanol/isopropanol (100:10, v/v). The gradient was 0–5 min = 5% B, 5–30 min = 90% B, 30–35 min = 90% B, and 35–45 min = 5% B, with a 5 μL injection volume in positive mode. LC-MS data were exported as mzData files using MassHunter Workstation [33].

#### Targeted Lipidomics Experiment

The targeted lipidomics experimental methods were described in detail in our earlier publication [34].

##### Lipid extraction

200 μL of water and 200 μL of methanol was added to each adipose tissue sample. 100 pmol of AA-d_8_ (Cayman Chemical, Ann Arbor, MI) was added to each tube as an internal standard for the fatty acid analysis. 100 pmol of PC(18:1/16:0-d_31_), PE(18:1/16:0-d_31_), PG(18:1/16:0-d_31_), PS(18:1/16:0-d_31_), and PI(18:1/16:0-d_31_) (Avanti Polar Lipids,) were added to each tube as internal standard for the phospholipid analysis. 100 pmol of LTC_4_-d_5_ and LTB_4_-d_4_ and 50 pmol of 12(13)-DiHOME-d_4_, 13-HODE-d_4_, 12(13)-EpOME-d_4_, PGD_2_-d_4_, PGE_2_-d_4_, 6-keto-PGF_1α_-d_4_, PGF_2α_-d_4_, TxB_2_-d_4_, 5-HETE-d_8_, 12-HETE-d_8_, and 15-HETE-d_8_ (Cayman Chemical) were added to each tube as internal standards for the lipid mediator analysis. 400 μL of methyl tertbutyl ether (MTBE) was then added to each sample to extract the lipids. The samples were incubated for 15 min on ice before the organic and aqueous phases were separated by centrifugation at maximum speed and 4 °C for 10 min. The MTBE layer (upper) was transferred to a clean glass tube for drying. The aqueous layer was extracted once more with 800 μL of MTBE. All fractions were combined and dried under a stream of nitrogen gas at room temperature. Each sample was dissolved in 4 mL of 3:1 isopropanol:ethanol for all downstream analyses.

##### AMP+ derivatization of fatty acids

An aliquot of 5 μL from each sample was placed into an LC-MS/MS vial insert and dried under a gentle nitrogen stream at room temperature. Samples were subsequently derivatized using the AMP+ protocol according to the manufacturer’s instructions (AMP+ MaxSpec Kit, Cayman Chemical) [35]. A derivatized fatty acid calibration mixture was prepared in parallel to verify the retention times of the corresponding derivatized fatty acids. In brief, the derivatization reagents supplied with the kit were added to each sample, followed by incubation at 60 °C for 30 min as specified in the AMP+ derivatization procedure. After cooling to room temperature, samples were subjected to LC-MS/MS analysis.

##### Protein isolation

After extraction, proteins present in the aqueous fraction were precipitated by adding 1 mL of ice-cold acetone. Samples were maintained on ice for 1 h to promote protein precipitation, followed by centrifugation at maximum speed for 10 min at 4 °C to pellet the proteins. The resulting protein pellets were resuspended in 300 μL of 1% SDS prepared in PBS and incubated at 65 °C for 1 h to ensure complete solubilization. Protein concentrations were then determined using a BCA assay according to the manufacturer’s protocol.

##### LC-MS/MS analysis of fatty acids

Samples were analyzed using a Shimadzu 8060NX triple quadrupole mass spectrometer coupled to a Shimadzu Nexera XS 40 series UHPLC system equipped with a Shimadzu CTO-40S column oven. A volume of 1 μL from each sample, blank, or standard mixture was injected and analyzed in positive ionization mode. Chromatographic separation was performed on a Shimadzu Nexcol C_18_ column (50 mm × 2.1 mm, 1.8 μm particle size, 100 Å pore size) fitted with a Phenomenex Security Guard C_18_ guard column (2.1 mm internal diameter). Mobile phase A consisted of 0.1% formic acid in water, while mobile phase B contained 0.1% formic acid in acetonitrile. The column temperature was maintained at 40 °C and the flow rate was held constant at 0.4 mL/min throughout the run. Initial conditions began at 20% mobile phase B for 0.5 min, followed by a linear gradient to 99% B over 4.5 min. The column was subsequently washed with 99% B for 2 min and re-equilibrated at 20% B for 5 min before the next injection. Instrument parameters and voltages were empirically optimized prior to analysis.

Multiple reaction monitoring (MRM) transitions for AMP+-derivatized fatty acids were as follows: m/z 471.3 → 183.1 for arachidonic acid, 479.3 → 183.1 for arachidonic acid-d_8_, 447.3 → 183.1 for linoleic acid, 473.3 → 183.1 for dihomo-γ-linolenic acid, 449.3 → 183.1 for oleic acid, 469.3 → 183.1 for eicosapentaenoic acid, 495.3 → 183.1 for docosahexaenoic acid, and 445.3 → 183.1 for γ-linolenic acid. Although certain fatty acids shared identical m/z transitions, chromatographic conditions enabled baseline separation of the corresponding peaks. Quantification was performed using calibration curves generated from AMP+-derivatized fatty acid standards. Peak areas from each analyte were normalized to the arachidonic acid-d8 internal standard to obtain absolute concentrations, followed by normalization to total protein content to account for differences in starting tissue mass.

##### LC-MS/MS analysis of phospholipids

Samples were analyzed using a Shimadzu 8060NX triple quadrupole mass spectrometer coupled to a Shimadzu Nexera XS 40 series UHPLC system and a Shimadzu CTO-40S column oven. A 5 μL aliquot of each sample or blank was injected and analyzed in both positive and negative ionization modes. Lipid separation was carried out on a Phenomenex Kinetex C_8_ column (150 mm × 2.1 mm, 2.6 μm particle size, 100 Å pore size) equipped with a Phenomenex Security Guard C_8_ guard column (2.1 mm internal diameter). Mobile phase A consisted of 5 mM ammonium formate in water, whereas mobile phase B was a 1:1 mixture of acetonitrile and isopropanol. The column temperature was maintained at 45 °C, and the flow rate was set to 0.3 mL/min throughout the analysis. The gradient program began at 20% mobile phase B and was held for 1 min, followed by a linear increase to 40% B over 1 min. Mobile phase B was then increased to 92.5% over the subsequent 23 min using a B-curve value of −3, followed by a linear increase to 98% B over 1 min. The column was washed at 98% B for 9 min and then re-equilibrated at 20% B for 8 min prior to the next injection. Instrument voltages and acquisition parameters were empirically optimized before sample analysis.

The Q1 m/z, Q3 m/z, and ionization mode corresponding to each analyte are provided in Supplemental Table S1. Quantification of phospholipids was performed by comparing analyte peak areas to those of the corresponding deuterated phospholipid internal standards within each headgroup class. Absolute lipid abundances were subsequently normalized to total protein content to account for differences in starting tissue material.

##### LC-MS/MS analysis of lipid mediators

Samples were analyzed using a Shimadzu 8060NX triple quadrupole mass spectrometer coupled to a Shimadzu Nexera XS 40 series UHPLC system and a Shimadzu CTO-40S column oven. A 5 μL volume of each sample or blank was injected and analyzed in both positive and negative ionization modes. Chromatographic separation was performed on a Phenomenex Kinetex C_8_ column (150 mm × 2.1 mm, 2.6 μm particle size, 100 Å pore size) fitted with a Phenomenex Security Guard C_8_ guard column (2.1 mm internal diameter). Mobile phase A consisted of 0.1% formic acid in water, while mobile phase B contained 0.1% formic acid in acetonitrile. The column oven temperature was maintained at 40 °C, and the flow rate was held constant at 0.4 mL/min throughout the run. The gradient began at 10% mobile phase B and increased to 25% B over the first 5 min, followed by a ramp to 35% B during the subsequent 5 min. Mobile phase B was then increased to 75% over the next 10 min. After separation, the column was washed with 98% B for 8 min and subsequently re-equilibrated at 10% B for an additional 8 min before the next injection. Instrument voltages and acquisition settings were empirically optimized prior to sample analysis. The Q1 m/z, Q3 m/z, and ionization polarity for each analyte are listed in Supplemental Table S2. Quantification was achieved by comparing the peak area of each analyte with that of the corresponding deuterated lipid mediator internal standard to obtain absolute concentrations. Lipid mediator levels were then normalized to total protein content to account for variations in the amount of starting tissue material.

#### Lipidomics data analysis

LINT-web was used for the analysis of lipidomics data [36]. The lipid expression data in .csv format was uploaded to the LINT-web tool. Under the data preprocessing step, lipid species with more than 70% missing values were removed before the analysis. In order to remove the variance between the samples, we utilized the median norm along with the log transformation and autoscaling. The Orthogonal partial least squares discriminant analysis (OPLS-DA) and principal component analysis (PCA) were studied to perform the dimensionality reduction on the samples. The differential expression of the lipids was studied using a Student’s t-test. The results were exported in various forms of graphs, like scores plot, heatmap, and volcano plot of lipid differences between the WT, APP, APP Cit, and APP Nit groups.

We calculated the elongation index (EI) and desaturation index (DI) to assess changes in fatty acid chain length and degree of unsaturation across lipid classes. For each lipid species, the total number of carbon atoms and double bonds was extracted from the lipid species (e.g., PC 36:2 corresponds to 36 carbons and 2 double bonds). For each lipid class (PC, PE, PS, PG, PI), lipid abundances were first averaged across replicates within each group. These indices were calculated separately for each comparison and used for downstream statistical analysis and visualization.

EI measures chain length progression within each lipid class (e.g., PC (28:1) → PC (30:1)):

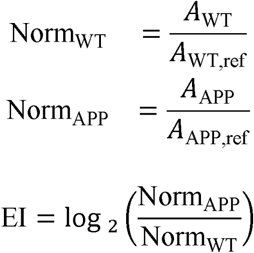

where ref = first lipid per class after sorting by unsaturation first, then chain length; A = averaged lipid abundance. DI measures unsaturation increases within fixed chain length blocks (e.g., PC (28:1) → PC (28:2)):

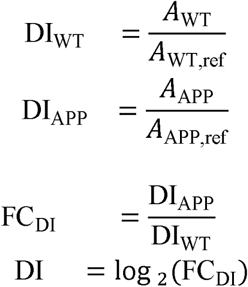

where ref = lowest unsaturation lipid per class within the same carbon chain block (e.g. PC (28:0), PC (28:1), PC (28:2)).

The circos plot of lipidomic changes was generated using the circlize R package [37, 38]. For the bar plot, the log of elongation index (log_2_ EI) and the log of desaturation index (log_2_ DI) were calculated for the three comparisons (APP vs WT, APP Cit vs APP, and APP Nit vs APP). The mean log EI and log DI values were summarized with 95% confidence intervals, and statistical significance was assessed using one-sample t-tests against zero. Significance was denoted as p < 0.05 (*), p < 0.01 (**), and p < 0.001 (***), while non-significant results were labeled ns.

#### BioPAN Pathway analysis

The lipid expression data in .csv format was submitted to the BioPAN pathway analysis tool [39], which was designed specifically for the lipidomics data (https://www.lipidmaps.org/biopan/). BioPAN determines statistical scores for all potential lipid pathways and their reactions, which may be active or suppressed. BioPAN uses the Z-score as a statistical measure that considers both the standard deviation and the mean to estimate the normal distribution of lipidomics data and identify significant pathways or lipid reactions (when *p* < 0.05).

### 2.6. Fluorescence Lifetime Imaging Microscopy of mice gWAT tissues

FLIM measurements were acquired using a Leica TCS SP8 confocal microscope equipped with motorized differential interference contrast (DIC) accessories. Imaging was performed with a white-light laser using an excitation wavelength of 525 nm and an emission detection range of 535–567 nm through a 20×/0.75 NA objective lens. FLIM images were collected with a pixel size of 568.18 nm × 568.18 nm and a pixel dwell time of 600 ns. Each image covered an area of 581.25 µm × 581.25 µm, and tandem scanner operation at 400 Hz corresponded to a frame rate shift of 0.024 s□¹. Fluorescence decay fitting parameters were optimized to obtain χ² values approaching 1, ensuring accurate lifetime fitting. [33].

### 2.7. Quantitative Reverse Transcription Polymerase Chain Reaction (qRT-PCR)

Total RNA from gWAT of different mouse groups was extracted using TRIzol reagent (Invitrogen) and reverse-transcribed into cDNA using MMLV reverse transcriptase and Oligo-dT primers (TaKaRa). qRT-PCR was performed with an UltraSYBR mixture (CWBIO) following the manufacturer’s instructions. Primers used are listed in Table S3. The GAPDH gene served as an internal reference. Ct values were normalized to β-actin, and relative differences between control and test groups were calculated using the 2^−ΔΔCt^ method, expressed as fold changes.

### 2.8. Raman Imaging of gWAT tissues

Raman spectra and imaging of gonadal white adipose tissue (gWAT) were acquired using a Renishaw inVia Reflex Raman spectroscopy platform equipped with a 50× long working distance objective lens. Spectra were collected using a 785 nm excitation laser with an exposure time of 20 s over a spectral range of 200–3200 cm□¹. Raman mapping was performed using the Map Image Acquisition mode. Spectral mapping parameters included a 1200 lines/mm grating, a 2 s integration time per pixel, and a spectral acquisition centre window of 1200 cm□¹.

### 2.9. Statistical Analysis

The data normality was evaluated using the D’Agostino–Pearson normality test. Differences among groups were determined by one-way analysis of variance (ANOVA) followed by Tukey’s multiple-comparison post hoc test. For energy expenditure analyses, ANOVA was performed with total body mass included as a covariate. Associations between variables were assessed using simple linear regression analysis. Principal component analysis (PCA) was carried out using Origin software (OriginLab, MA, USA). Results are presented as mean ± standard deviation (SD). Statistical analyses were conducted using GraphPad Prism v10.2.0 (GraphPad Software, Boston, MA, USA). A p value < 0.05 was considered statistically significant.

## 3. Results

### 3.1. APP/PS1 mice displayed impaired lipolysis

APP/PS1 mice maintained on a high-fat diet exhibited a significant increase in body mass (**Fig. 1a**) and body weight (**Fig. 1b, 1c**) compared with WT controls, indicating elevated systemic metabolic burden. Neither citrulline nor nitrite treatment significantly altered body mass or adiposity relative to untreated APP mice. Consistent with these findings, **Fig. 1f** showed that adipocyte size was significantly increased in gWAT from APP, APP mice treated with citrulline, and nitrite groups compared with WT, demonstrating persistent adipocyte hypertrophy despite intervention. Ex vivo lipolysis analysis of gWAT explants revealed a significant reduction in glycerol release in APP, APP treated with citrulline, and nitrite mice relative to WT (**Fig. 1d)**, indicating impaired lipolytic capacity and reduced triglyceride breakdown across all APP groups. Our, glucose tolerance testing (**Fig. 1e**) showed significantly elevated glucose area under the curve (AUC) in APP mice at both 15 and 25 weeks compared with WT, consistent with glucose intolerance. Citrulline treatment modestly reduced glucose AUC at 25 weeks relative to APP mice, whereas nitrite treatment did not produce a significant improvement. Together, these results demonstrate that APP/PS1 mice develop obesity-associated adipocyte hypertrophy, impaired lipolysis, and glucose intolerance that are not reversed by NO-directed interventions, although citrulline treatment to led a modest improvement in glucose handling at 25 week.

**Figure 1.**
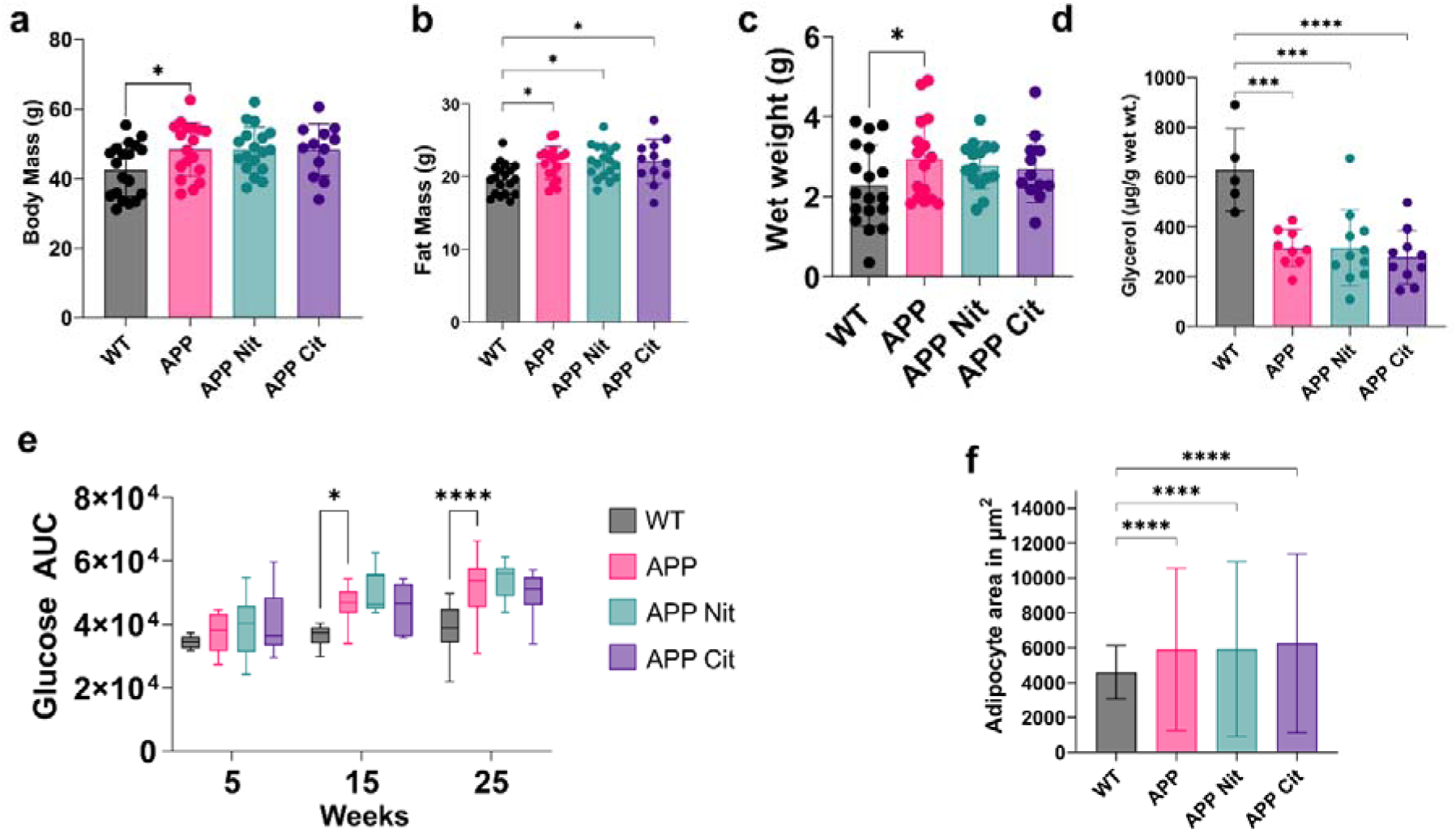
Body mass, adiposity, and metabolic tolerance in WT and APP/PS1 mice subjected to a high-fat diet (HFD) with nitrite or citrulline treatment. (a) Longitudinal changes in body mass over time. Comparison of (b) fat mass and (c) gonadal white adipose tissue (gWAT) wet weight among HFD-fed WT (Control), APP, APP + nitrite (APP Nit), and APP + citrulline (APP Cit) mice. (d) Ex vivo glycerol release from gWAT explants as a measure of lipolytic activity. (e) Glucose tolerance test (GTT) and corresponding area under the curve (AUC) for each group a 5, 15, and 25 weeks of treatment. (f) Quantification of adipocyte area across experimental groups.

### 3.2 APP/PS1 mice exhibit impaired phospholipid turnover

Comprehensive targeted lipidomic analysis of gWAT revealed broad and consistent alterations in phospholipid composition in APP/PS1 mice maintained on a high-fat diet. Fold changes (FC) were calculated for APP versus WT, APP + nitrite versus APP, and APP + citrulline versus APP. In the heatmaps, green indicates FC < 1 (decrease), white indicates FC = 1 (no change), and red indicates FC > 1 (increase). Compared with WT controls, APP mice exhibited marked downregulation of lysophospholipids and major membrane phospholipids across nearly all species analyzed. Lysophosphatidylcholines (LPCs) were uniformly reduced in APP mice, with most LPC species displaying fold changes (FC) in the range of approximately 0.5–0.8 relative to WT (**Fig. 2b**). A similar pattern was observed for phosphatidylcholines (PCs) (**Fig. 2a**), Lysophosphatidylethanolamines (LPEs) (**Fig. 2c**), and phosphatidylethanolamines (PEs) (**Fig. 2d**), with the majority of species showing FC (APP/WT) values between 0.4 and 0.8, indicating a global suppression of membrane phospholipid abundance in APP adipose tissue. Evaluation of treatment effects relative to untreated APP mice revealed distinct lipidomic signatures for nitrite and citrulline. In the LPC class, nitrite treatment produced partial restoration, with most LPC species exhibiting FC (APP Nit/APP) values between approximately 0.7 and 0.9, whereas citrulline treatment resulted in stronger recovery for several LPC species, with FC (APP Cit/APP) approaching or exceeding unity and reaching up to ∼1.3 for select species (Fig. 2b). PC species showed modest rescue with nitrite treatment, particularly among shorter-chain PCs, where FC (APP Nit/APP) values were in the range of ∼1.0–1.3 (Fig. 2a). Citrulline also partially restored PC levels, though the magnitude of rescue was more variable across species. Phosphatidylethanolamine (PE) species, including several long-chain and polyunsaturated species associated with mitochondrial membranes, demonstrated more pronounced recovery with treatment. Both nitrite and citrulline increased multiple PE species relative to APP, with citrulline producing larger increases for several species, including PE(40:1), PE(42:3), and PE(42:7), for which FC (APP Cit/APP) values ranged from approximately 1.3 to 1.6 (Fig. 2d). Phosphatidylglycerol (PG), a precursor for cardiolipin and a critical component of mitochondrial membranes, exhibited a mixed response overall; however, several PG species were strongly increased with treatment. Notably, PG(34:0) showed a dramatic increase with citrulline treatment relative to APP, with FC (APP Cit/APP) approaching 49-fold, while PG(36:3) and PG(42:7) were also elevated following treatment (**Fig. 2f**).

**Figure 2.**
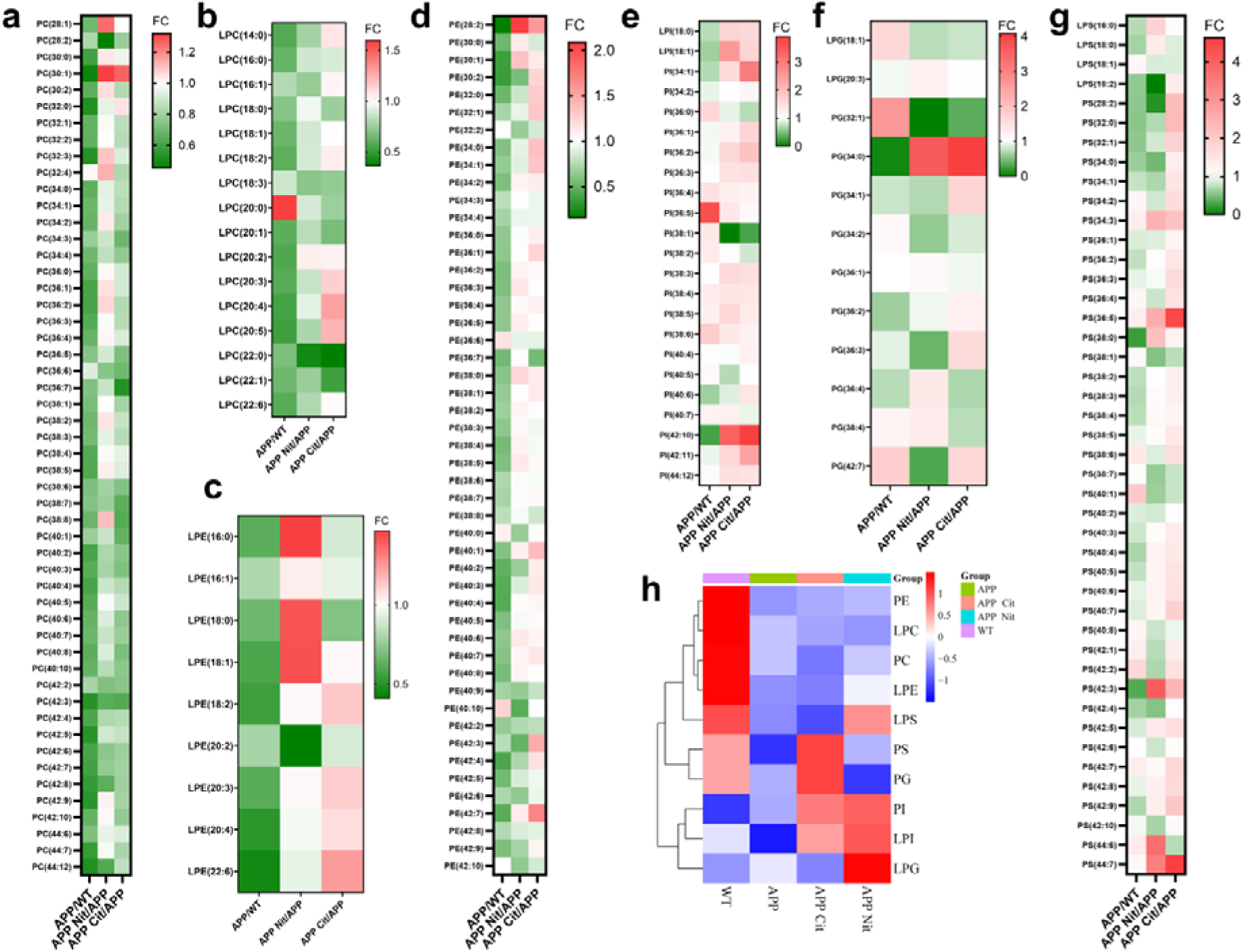
Targeted phospholipid remodeling in gWAT of WT and APP/PS1 mice and the effects of nitrite and citrulline treatment. (a–g) Heatmaps depicting fold changes (FC) of individual lipid subspecies within major phospholipid and lysophospholipid classes, including (a) phosphatidylcholines (PC); (b) lysophosphatidylcholines (LPC); (c) lysophosphatidylethanolamines (LPE); (d) phosphatidylethanolamines (PE); (e) phosphatidylinositols (PI & LPI); (f) phosphatidylglycerols (PG & LPG); (g) phosphatidylserines (PS & LPS). Fold changes were calculated for APP versus WT, APP + nitrite versus APP (APP Nit/APP), and APP + citrulline versus APP (APP Cit/APP). Green indicates FC < 1 (decrease), white indicates FC = 1 (no change), and red indicates FC > 1 (increase). (h) Heatmap showing class-level changes based on the averaged sum of all detected subspecies within each phospholipid (PL) class. Z-scores were calculated from log2-transformed lipid intensity values.

Phosphatidylinositol (PI) species displayed heterogeneous changes in APP mice but were broadly increased following both nitrite and citrulline treatment. Nitrite treatment produced particularly robust increases in several PI species, while citrulline also elevated multiple PI lipids, with FC (APP Cit/APP) values ranging from approximately 1.3 to nearly 4.0 for select species (**Fig. 2e**). In contrast, phosphatidylserine (PS) species, lipids commonly associated with membrane stress and apoptotic signalling, were elevated in APP mice relative to WT and increased further following treatment. Nitrite treatment induced particularly large increases in several PS species, ranging from 2–5-fold relative to APP. Although citrulline treatment also increased PS levels, the increase was modest (**Fig. 2g**). From our results, these lipidomic data demonstrate that APP/PS1 mice on HFD exhibit widespread suppression of lysophospholipids and membrane phospholipids in adipose tissue. Both nitrite and citrulline shift the lipidome toward WT values, but with distinct class-specific patterns: nitrite preferentially enhances signaling-associated lipids such as PI and PS, whereas citrulline produces broader recovery of LPC, PC, PE, and mitochondrial-associated PG species, yielding a more balanced phospholipid remodeling profile in gWAT. **Figure 2h** shows the heatmap of the averaged summed intensities of all detected subspecies within each phospholipid (PL) class. The heatmap is generated using Z-scores calculated from the log2-transformed lipid intensity values for all species. The results demonstrate broad upregulation of lysophospholipids (LPC, LPE, and LPS) and phospholipids (PC, PE, PS, and PG) in the WT group, whereas LPG, LPI, and PI were downregulated in WT. In contrast, all phospholipid classes were downregulated in APP mice. Citrulline treatment (APP Cit) resulted in upregulation of PS, PG, PI, and LPI, while nitrite treatment (APP Nit) led to recovery of LPS, LPI, LPG, and PI, which were upregulated relative to APP.

### 3.3 Ceramide and TG enrichment in APP/PS1 adipose tissues

Untargeted lipidomic profiling of gWAT revealed distinct global lipid signatures among WT, APP, APP + citrulline (APP Cit), and APP + nitrite (APP Nit) groups. Partial least squares–discriminant analysis (PLS-DA) demonstrated clear separation of all four groups, with component 1 and component 2 accounting for 18.9% and 17.8% of the total variance, respectively (**Fig. 3a**). Variable importance in projection (VIP) analysis identified multiple lipid species with VIP scores >1, indicating strong contributions to group discrimination and highlighting ceramides, triacylglycerols (TG), monoacylglycerols (MG), and phosphatidic acids (PA) as key differentiating lipid classes (**Fig. 3b**). Class-level heatmap analysis based on Z-scores of log2-transformed lipid intensities showed that WT mice exhibited relative downregulation of TG, PG, ceramides, CDP-diacylglycerol (CDP-DG), PA, and MG, whereas APP mice displayed broad upregulation across these lipid classes (**Fig. 3c**). Relative to APP, both citrulline and nitrite treatment resulted in decreased abundance of most lipid classes; however, the magnitude of reduction was greater in APP Cit than in APP Nit. Notably, MG levels were increased in APP Cit compared with APP, whereas PA levels were markedly reduced, while APP Nit exhibited a significant decrease in MG relative to both APP and APP Cit.

**Figure 3.**
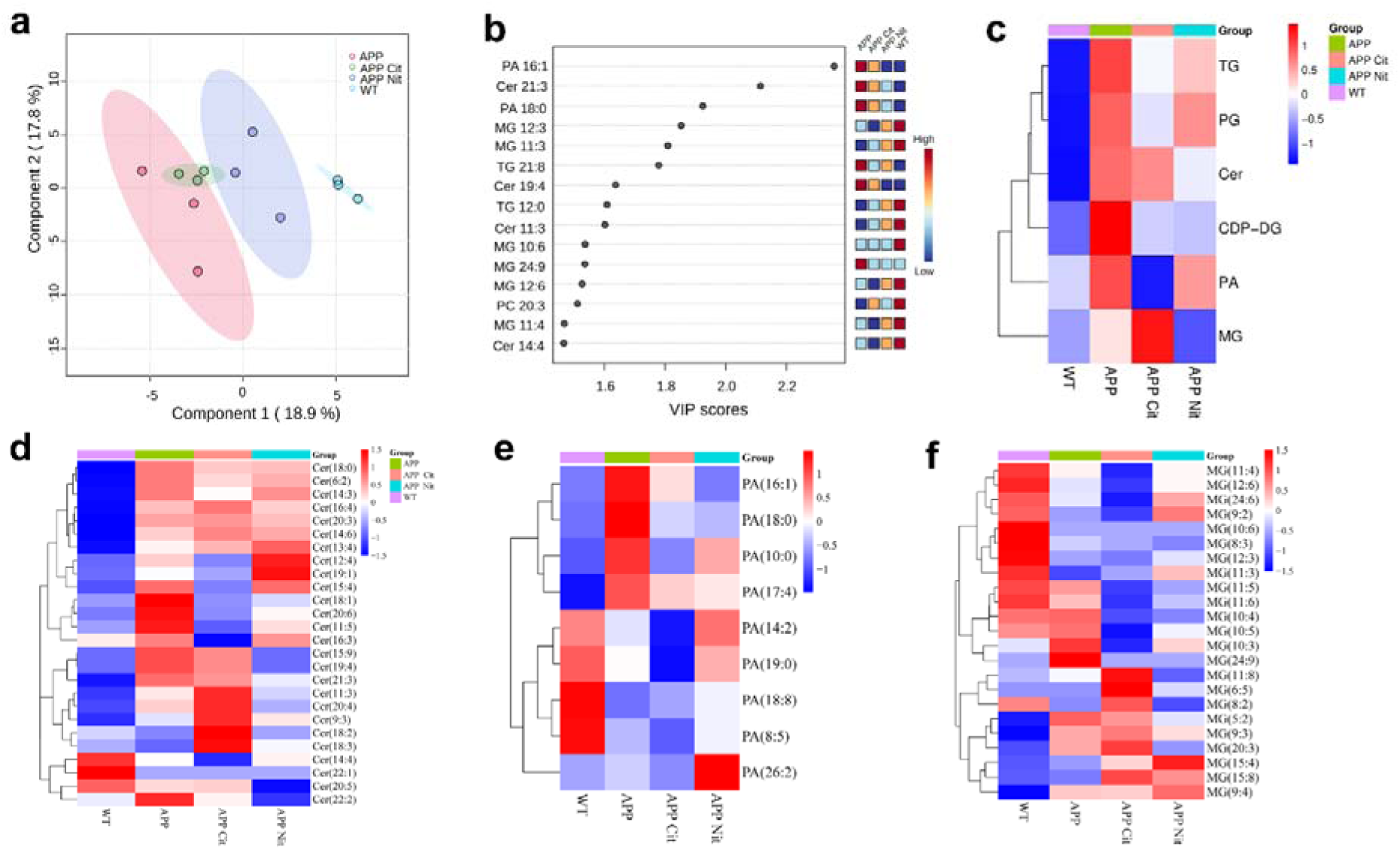
Untargeted lipidomic profiling of gWAT of APP/PS1 mice and differential modulation by citrulline and nitrite. (a) Partial least squares discriminant analysis (PLS-DA) score plot showing distinct clustering of WT, APP, APP + citrulline (APP Cit), and APP + nitrite (APP Nit) groups. Components 1 and 2 account for 18.9% and 17.8% of the total variance, respectively. (b) Variable importance in projection (VIP) scores for differentially abundant lipid species, highlighting lipids with VIP > 1 as major contributors to group separation. (c) Class-level heatmap showing changes in major lipid classes based on the averaged sum of all detected subspecies within each class. Z-scores were calculated from log2-transformed lipid intensity values. (d–f) Heatmaps of species-level changes for ceramides (d), phosphatidic acids (e), and monoacylglycerols (f), illustrating lipid class–specific responses to APP genotype and treatment.

At the species level, ceramides were consistently elevated in APP mice compared with WT, with most species increased by approximately 1.05–1.4 fold, indicating ceramide accumulation in APP adipose tissue (**Fig. 3d**). Citrulline and nitrite treatment produced only modest reductions in ceramide levels relative to APP, with most species remaining near baseline. TG species displayed marked heterogeneity, with many TGs moderately increased in APP versus WT and select species showing pronounced elevations following citrulline or nitrite treatment (e.g., TG(23:4) exhibiting a >5-fold increase in APP Cit compared with WT) (Fig. 3c). MG species, intermediates of lipolysis, were generally mildly increased in APP compared with WT, with exceptions such as MG(12:3), which was markedly reduced. Both treatments increased several MG species relative to WT, and MG levels were higher in APP Cit compared with APP. APP Nit exhibited a reduction in MG abundance relative to both APP and APP Cit (**Fig. 3f**). PA species showed large and variable changes across groups, with several PA species elevated in APP relative to WT and selectively increased with citrulline treatment, while responses to nitrite were more limited (**Fig. 3e**). Together, the untargeted lipidomic data demonstrate that APP/PS1 mice on high-fat diet exhibit widespread accumulation of storage and bioactive lipid classes, including ceramides, TGs, MGs, and PA. Further, citrulline and nitrite differentially modulate these lipid pools, with citrulline producing broader class-level normalization and nitrite eliciting a more modest reduction relative to APP.

### 3.2. Pro -inflammatory lipid mediators are downregulated after sodium nitrite and L-citrulline treatment

We performed analysis to evaluate the metabolic shifts in fatty acid chain length (Elongation Index) and double bond density (Desaturation Index) across different groups (**Fig. 4**). The results demonstrate that the APP condition is associated with a clear and consistent increase in lipid elongation compared to WT. Figure 4b showed a significant increase in the elongation index across multiple lipid classes, including PE, PC, PS, and PG (**Figs. 4b-4g**). In contrast, desaturation changes are minimal and largely non-significant overall, although class-specific differences are observed, with PC showing a significant decrease in desaturation (**Fig. 4d**) and PI exhibiting reduced elongation (**Fig. 4g**). The circos plot (**Fig. 4a**) further supports these findings, highlighting a predominance of longer chain lipid species and several significantly altered lipids in APP, indicating that disease-associated lipid remodeling is driven primarily by enhanced fatty acid elongation.

**Figure 4.**
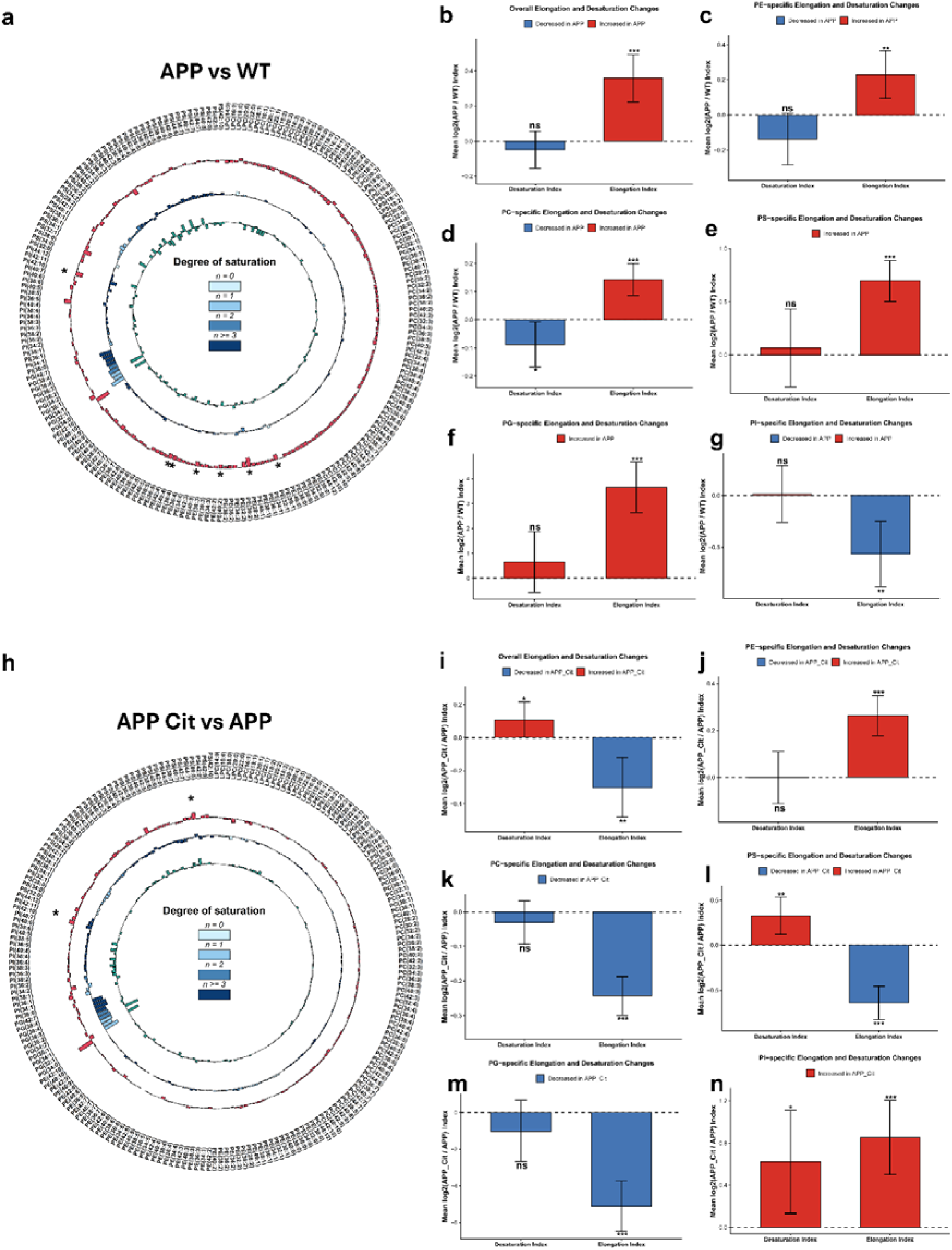
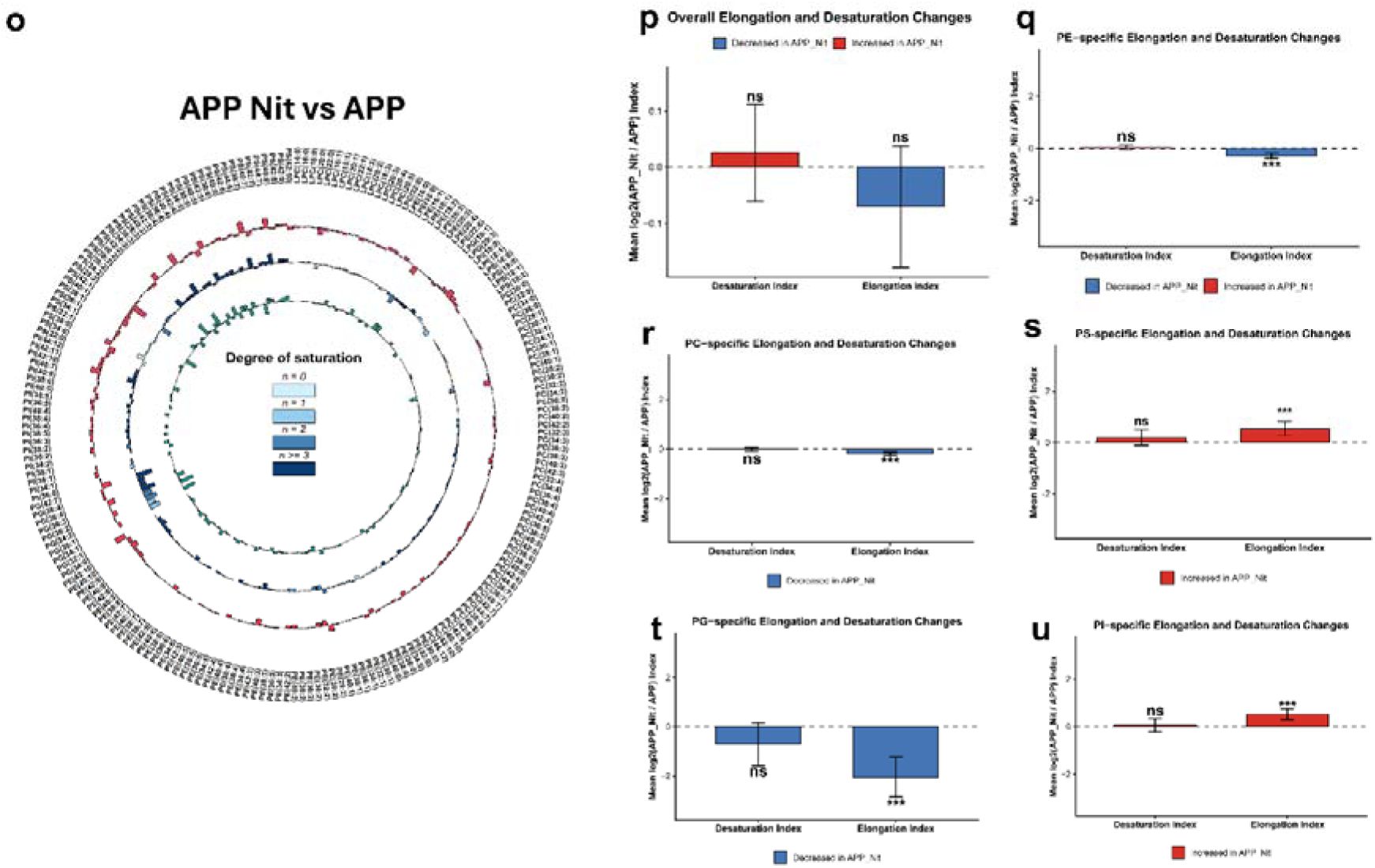
Impact of nitrite and citrulline treatments on global and class-specific lipid desaturation and elongation indices. (a) APP vs. WT: Circos plots visualize differential lipid species abundance categorized by degree of unsaturation (n=0 to n≥3). Bar graphs represent the mean log2 fold-change of Desaturation and Elongation indices for total lipids and specific phospholipid classes: (b) overall, (c) Phosphatidylethanolamine (PE), (d) Phosphatidylcholine (PC), (e) Phosphatidylserine (PS), (f) Phosphatidylglycerol (PG), and (g) Phosphatidylinositol (PI). APP mice exhibit a significant systemic increase in the Elongation Index across most lipid classes compared to WT. (h-n) APP Cit vs. APP: Comparison of Citrate-treated APP mice against untreated APP controls. Citrate treatment appears to significantly reverse the elongation trends observed in the APP model, particularly in PC, PS, and PG classes, while showing a modest increase in the Desaturation Index for total lipids. (o-u) APP Nit vs. APP: Comparison of Nitrate-treated APP mice against untreated APP controls. Nitrate treatment results in a significant reduction of the Elongation Index across several classes (PE, PC, PG) without significantly altering the global Desaturation Index. Statistical significance was determined via t-test]; ∗p<0.05, ∗∗p<0.01, ∗∗∗p<0.001; ns, non-significant.

Upon treatment, citrulline markedly alters this pattern by significantly reducing the elongation index in APP (**Fig. 4i**), suggesting a reversal of the elongated lipid profile, while inducing a mild increase in desaturation. This effect is consistent across PC (**Fig. 4k**), PS (**Fig. 4l**), and PG (**Fig. 4m**) classes, whereas PE (**Fig. 4j**) shows increased elongation and PI (**Fig. 4n**) displays increases in both elongation and desaturation, indicating a class-specific response. The circos plot (**Fig. 4h**) supports these changes, showing a shift away from highly elongated lipid species. In contrast, nitrite treatment produces more limited and inconsistent effects with no significant global changes in elongation or desaturation indices. However, class-specific changes are evident, including a significant decrease in elongation in PC (**Fig. 4r**) and PG (**Fig. 4t**). Elongation increased significantly in PS (**Fig. 4s**) and PI (**Fig. 4u**), while desaturation remained largely unchanged. The circos plot (**Fig. 4o**) reflects this mixed pattern, lacking a clear global trend. Overall, the findings indicate that APP-associated lipid alterations are largely driven by enhanced fatty acid elongation. Citrulline treatment appears to partially restore this imbalance by suppressing elongation, while nitrite exerts more selective, class-specific effects and does not achieve clear overall normalization of the lipid profile.

Pathway analysis of the lipids (**Fig. S1a-1c**) for targeted lipidomics was performed using LipidOne. Blue arrows indicate reactions that are activated, and red arrows mark reactions that are suppressed. In the targeted APP vs WT lipidomics analysis (**Fig. S1a**), the lipid pathway shows specific phospholipid conversion steps. Blue arrows indicate reactions that are activated in APP, such as the conversion of PE 40:4 and PE 40:8 to their corresponding PS species via Ptdss1/2, leading to accumulation of long, unsaturated PS 36:4, PS 40:4, and PS 42:7 (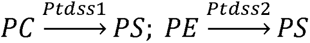. Red arrows mark reactions that are suppressed, including the back-conversion of PS to PE via Psd 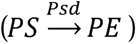 and the methylation of PE 32:2 to PC 32:2 via Pemt 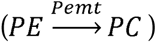, which results in lower flux toward PE regeneration and PC synthesis. In the targeted APP Cit vs WT lipidomics analysis (**Fig. S1b**) reactions catalysed by Ptdss1/2 that convert PE to PS are relatively activated. On the other hand, Psd□mediated PS→PE and Pemt□dependent PE→PC steps are suppressed, leading to reduced flux toward PC and partial accumulation of specific PS and PE species. This pattern, captured by positive Z□scores for Ptdss1/2 and strongly negative values for Psd and Pemt, suggests that citrulline shifts the APP samples toward a more PS enriched, less Pemt□driven membrane phospholipid profile. In APP Nit vs APP (**Fig. S1c**), the reactions catalysed by Ptdss1/2 that drive PE and PC toward PS 42:8 are strongly activated (positive Z-scores and blue arrows), while the reverse PS→PE reaction via Psd and one PE→PC 38:7 step via Pemt are inhibited (negative status, red arrows). This pattern suggests that nitrite drives metabolic flux toward building up highly unsaturated PS 42:8, while reducing its conversion back to PE or onward to PC. BioPAN pathway analysis of untargeted lipidomics (**Fig. S1d-S1fl**) compares PA and PC species between groups. For PA (10:0) to PC (10:0), there is strong upregulation in APP vs WT (**Fig. S1d**) (score 2.488), some upregulation in APP vs APP Nit (0.116) (**Fig. S1f)**, and moderate downregulation in APP vs APP Cit (−1.018) (**Fig. S1e)**. For PA (17:4) and PC (17:4), APP vs WT (Fig. S1d) and APP vs APP Nit (Fig. S1f) showed weak positive shifts (0.163, 0.531), while APP vs APP Cit (Fig. S1e) showed a notable negative shift (−1.381). PA (18:0) to PC (18:0) does not change in any comparison (score 0 throughout). Overall, this suggests specific phosphatidic acid to phosphatidylcholine conversions, that are strongly upregulated in APP compared to WT and APP Nit, but were suppressed in APP Cit.

These changes in fatty acid elongation and desaturation patterns are further supported by global lipidomic index analysis (Supplementary Figure S2). Citrulline and nitrite treatments significantly modulated elongation indices across phospholipid classes. Species-level analysis further revealed that APP mice exhibit coordinated changes in phospholipid chain length and unsaturation across PC, PE, PI, PS, and PG classes, as shown by hierarchical clustering and chain-length distribution patterns (Supplementary Figure S2a). Although long-to-short chain ratios showed an increasing trend in APP mice, these differences were not statistically significant across most classes, suggesting subtle but coordinated lipid remodeling (Supplementary Figure S2b–g). Receiver operating characteristic (ROC) analysis demonstrated that total phospholipids and PI species possess the highest discriminatory power between WT and APP groups (Supplementary Figure S2h). Citrulline treatment induced selective remodeling of phospholipid species, particularly within PE, PI, and PG classes, as shown by changes in species distribution and chain-length characteristics (Supplementary Figure S3a). Consistent with these findings, long-to-short chain ratios exhibited a trend toward normalization following citrulline treatment, particularly in PE and PI classes, although these changes did not reach statistical significance (Supplementary Figure S3b–g). ROC analysis further indicated that PE species provide the strongest discriminatory signature for citrulline treatment effects (AUC = 0.889), suggesting a class-specific metabolic response (Supplementary Figure S3h). Nitrite treatment resulted in distinct phospholipid remodeling patterns, particularly affecting chain-length distribution across lipid classes (Supplementary Figure S4a). A trend toward increased long-to-short chain ratios was observed in total phospholipids and PS classes following nitrite treatment, although these changes were not statistically significant (Supplementary Figure S4b–g). Notably, ROC analysis revealed that PS species exhibited perfect classification performance (AUC = 1.000), highlighting phosphatidylserine remodeling as a key signature of nitrite-mediated lipid regulation (Supplementary Figure S4h). To further evaluate the functional implications of lipid remodeling, we performed a lipid functional analysis (Supplementary Figure S5). APP mice exhibited a shift toward long-chain fatty acid enrichment, as indicated by increased short/long and long/medium chain ratios, suggesting altered energy metabolism and a preference for β-oxidation pathways. In parallel, structural indices demonstrated increased double bond index and reduced saturation index, indicating enhanced membrane unsaturation and fluidity. Notably, signaling-related indices revealed a marked increase in PI/PL ratio, consistent with activation of intracellular signaling pathways, while reductions in lysophospholipid indices (LPC+LPE)/PL and LPC/PC suggest decreased phospholipid turnover. Together, these findings indicate that APP-associated lipid remodeling extends beyond compositional changes to impact membrane structure, signaling dynamics, and metabolic function.

We further analyzed the lipid mediators using targted lipidomics approach. Our results showed that the oxidized lipid mediators 12-HETE (**Fig. 5b**), 13-HODE (**Fig. 5c**), and 14-HDHA (**Fig. 5d**) were significantly increased in APP mice compared with WT, indicating enhanced oxidative lipid signaling in the APP/HFD condition. 15-HETE (**Fig. 5a**) was also relatively higher in APP compared to WT. Citrulline- and nitrite-treated APP mice displayed lower levels of these mediators relative to untreated APP animals; however, these reductions did not reach statistical significance. Additional oxylipins and endocannabinoid-related mediators (9-HODE, 11-HETE, AEA, OEA) exhibited similar directional trends (see Supporting Information Figure S6) but showed no statistically significant differences among groups.

**Figure 5.**
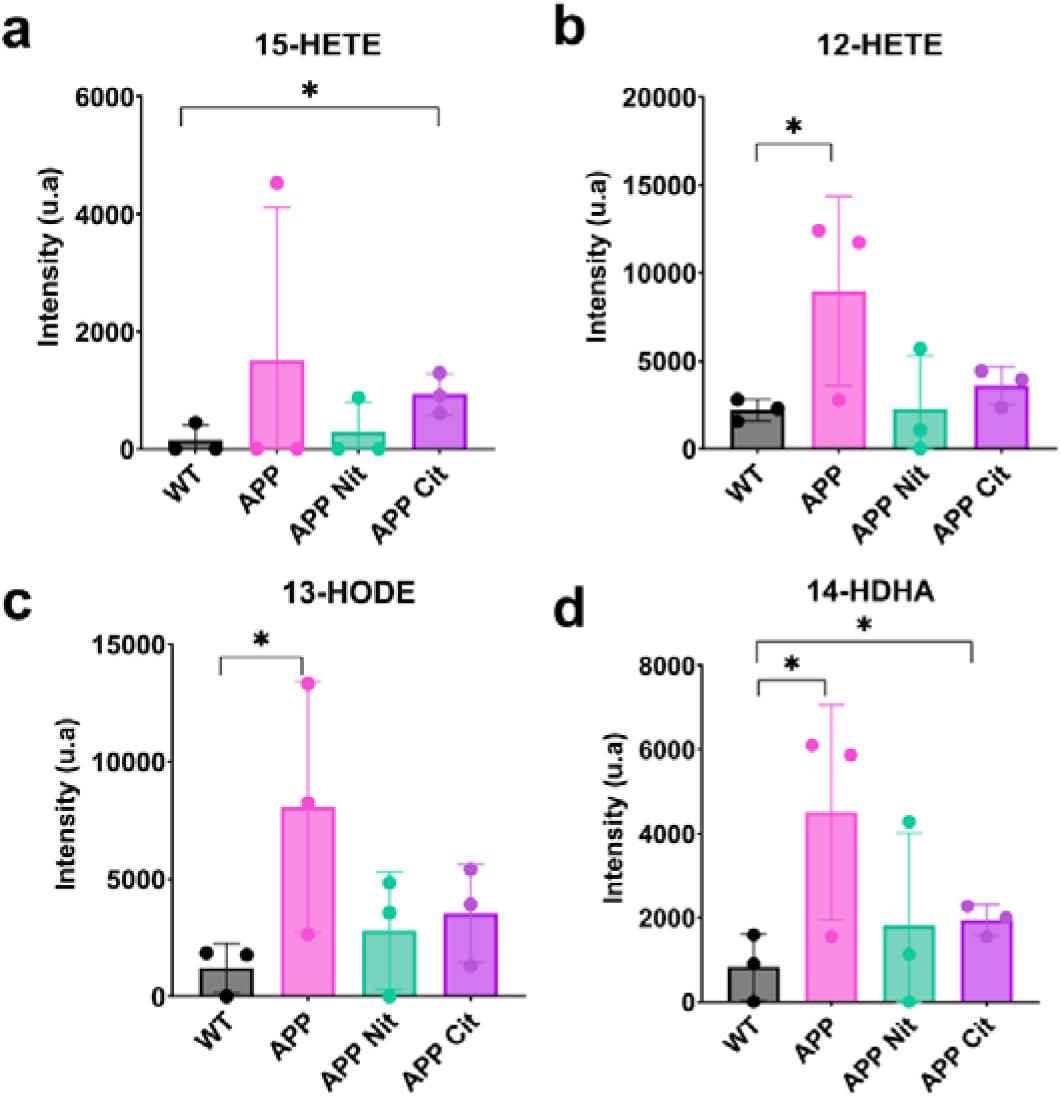
Pro-inflammatory lipid mediators are downregulated after Nitrite and Citruline treatment. Comparison of (a) 15-Hydroxyeicosatetraenoic acid (15-HETE), (b) 12-Hydroxyeicosatetraenoic acid, (c) 13-Hydroxyoctadecadienoic acid (13-HODE), and (d) 14-hydroxy-docosahexaenoic acid (14-HDHA) for WT, APP, APP Nit and APP Cit.

#### Raman microscopy mapping of nitro–fatty acids

To spatially map NO-derived lipid species in gWAT, we performed Raman microspectroscopy combined with spectral mapping of nitro–fatty acids (**Fig. 6a**). Nitro-oleic acid (nitro-OA) was significantly reduced in APP mice compared with WT (**Fig. 6d**). In contrast, APP mice exhibited significantly higher levels of nitro-DHA (**Fig. 6f**), nitro–conjugated linoleic acid (nitro-CLA) (**Fig. 6h**), and nitro-arachidonic acid (nitro-AA) (**Fig. 6j**) compared with WT. This indicates enhanced nitration of polyunsaturated fatty acids in APP fed HFD group under elevated oxidative and nitrative stress. Citrulline and nitrite treatment significantly increased nitro-OA relative to untreated APP mice and simultaneously reduced nitro-CLA, nitro-AA, and nitro-DHA back toward WT levels. These changes were evident in adipocyte regions and perivascular areas of gWAT, demonstrating that NO-directed interventions not only alter global lipid composition but also qualitatively remodel the pattern of nitro–fatty acid formation in situ. The non-nitrated corresponding fatty acids (OA, CLA, AA, and DHA) did not show differences among groups although the mean values of these fatty acids in APP were higher compared to WT groups (**Figs. 6c, 6e, 6g,** and **6i**). The Raman spectra of the nitro fatty acids are shown in **Fig. 6k**. The Raman spectra showed distinct peaks in the fingerprint region (200 – 2000 cm^−1^) as well as the high wavenumber (2500 – 3200 cm^−1^) region. The Raman peaks and their corresponding vibrational assignments are provided in Table S4 (see Supplementary Information). The corresponding principal component analysis (PCA) of these spectra showed distinct clusters (**Fig. 6l**), suggesting that the Raman spectra could be used as an optical marker for each nitro fatty acids.

**Figure 6.**
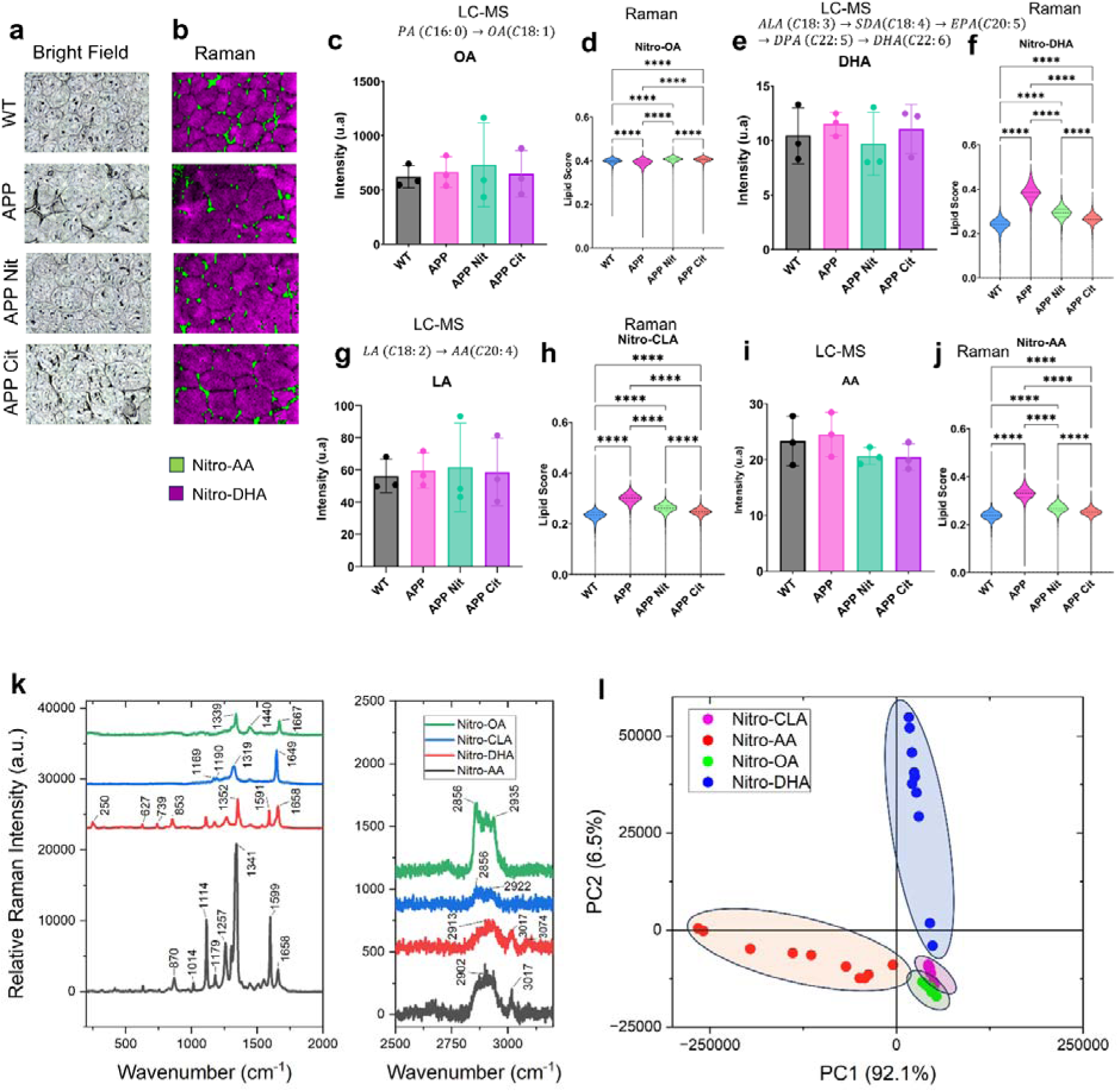
Raman spectroscopy and imaging reveal differential spatial distribution of nitro-fatty acids across experimental groups. Representative (a) brightfield images and (b) corresponding Raman maps from WT, APP, APP + nitrite (APP Nit), and APP + citrulline (APP Cit) mice illustrating the spatial distribution of nitro-arachidonic acid (NO□-AA) and nitro-docosahexaenoic acid (NO□-DHA). Quantitative comparisons of native fatty acids measured by LC–MS and their corresponding nitro-fatty acid species measured by Raman spectroscopy are shown for (c) oleic acid (OA), (d) nitro-oleic acid (NO□-OA), (e) docosahexaenoic acid (DHA), (f) NO□-DHA, (g) linoleic acid (LA), (h) nitro-conjugated linoleic acid (NO□-CLA), (i) arachidonic acid (AA), and (j) NO□-AA. (k) Representative Raman spectra comparing NO□-OA, NO□-CLA, NO□-DHA, and NO□-AA. (l) Principal component analysis (PCA) of Raman spectra demonstrating distinct clustering of individual nitro-fatty acid species.

To map the metabolic changes in the gWAT samples, we performed fluorescence lifetime imaging microscopy (FLIM). The FLIM images revealed a significant increase in mean fluorescence lifetime (τ_m_) in APP mice compared with WT (**Fig. 7d**), indicating a shift toward protein-bound NADH and impaired mitochondrial redox cycling. Both APP Cit and APP Nit groups also displayed elevated τ_m_ relative to WT (Fig. 7d), reflecting persistent mitochondrial stress despite treatment. **Figure 7a** shows the FLIM images of all four groups and **Fig. 7b** shows the corresponding polar plot for the lifetime images. The lifetime spectra are shown in **Fig. 7c**. These findings are consistent with the lipidomic signatures of decreased phospholipid remodeling, elevated ceramides, and increased oxidized lipid mediators in APP mice, all of which impair mitochondrial membrane organization and redox activation. The persistence of elevated τ_m_ in APP Nit matches with Nox4 upregulation in the RT-PCR data (Fig. 9), indicating redox activation. While the partial lipid rescue observed in APP Cit suggests improved but not fully restored mitochondrial metabolism.

**Figure 7.**
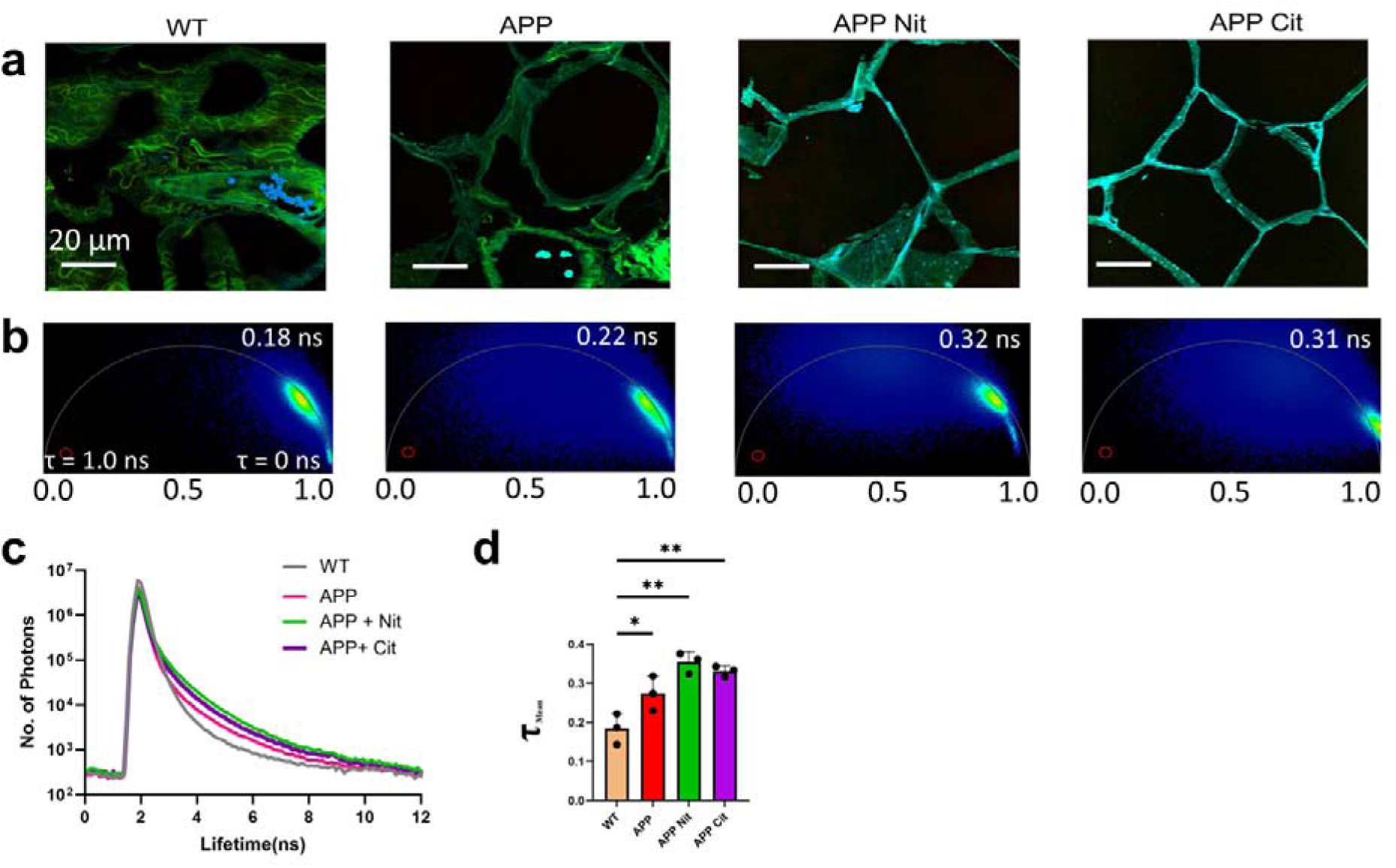
Mapping metabolic changes in the gWAT using fluorescence lifetime imaging microscopy (FLIM). (a) FLIM images for different groups (WT, APP, APP Nit, APP Cit). (b) The phasor plot for the different groups showing that the lifetimes are multicomponent. (c) Comparison of mean lifetime spectra from different groups. (d) Comparison of mean lifetime for different groups.

To better understand the effects of our treatments, we examined the brains of mice from each experimental group, focusing on the hippocampus, a key brain region involved in learning, memory, and spatial awareness. Within the hippocampus, the cornu ammonis (CA1–CA3) subfields and the dentate gyrus (DG) play unique roles in maintaining brain function and are often affected in AD [40, 41]. From the H&E staining (**Fig. 8**), we assessed the overall hippocampal architecture. In WT mice fed an HFD, we observed well-preserved brain architecture, with densely packed pyramidal neurons in CA1 and CA3 and clearly defined granule cells in the DG. In contrast, APP mice showed clear signs of disrupted neuronal organization, loss of structure, and cell death, particularly in the CA1 and CA3 areas. Interestingly, the brains of APP Nit and APP Cit mice showed much better preservation of hippocampal structure. Their neuronal layers looked more intact, and there were fewer signs of degeneration, suggesting that both treatments may have a protective effect on the brain.

**Figure 8.**
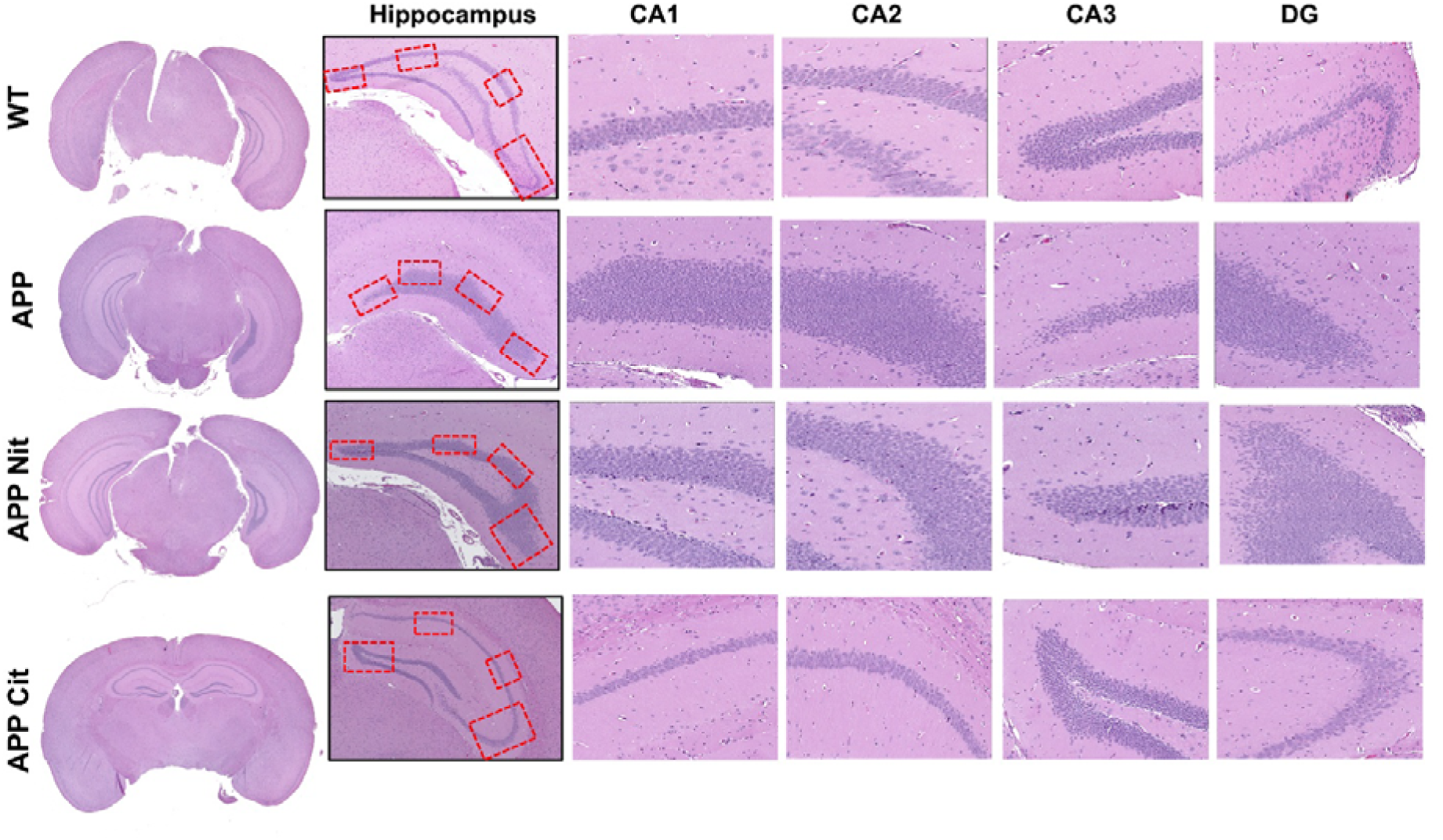
Sodium nitrite and citrulline protect neuronal cells in the hippocampus of mouse brains. APP/PS1 mice were treated with HFD and in combination with either Sodium nitrite or citrulline. H&E staining images from the WT, APP, APP Nit, and APP Cit groups. The hippocampus and CA1, CA2, CA3, and DG regions of the hippocampus are shown for each group.

Analysis of inflammatory, redox, and nitric oxide–related gene expression in gonadal white adipose tissue revealed broad activation of oxidative and inflammatory pathways in APP/PS1 mice (**Fig. 9e–k**). RT-PCR analysis showed that Nox4 expression was significantly increased in APP (*p* < 0.05), APP + citrulline (APP Cit; *p* < 0.01), and APP + nitrite (APP Nit; *p* < 0.01) compared with WT, with APP Cit exhibiting a smaller increase relative to APP Nit (**Fig. 9h**). Tnf-α mRNA was significantly elevated in APP (*p* < 0.05) and APP Cit (*p* < 0.01) compared with WT, while APP Nit showed a non-significant increase (**Fig. 9j**). Nos2 expression did not differ among groups, whereas Nos3, Catalase (Cat), thioredoxin 1 (Txn1), and Il6 were increased in APP, APP Nit, and APP Cit relative to WT, reaching statistical significance only for Txn1 in APP Nit (**Fig. 9e–g, 9k**). Sod1 expression was significantly increased in APP Nit compared with APP, while APP Cit showed intermediate levels that were not significantly different from WT or APP Nit (**Fig. 9i**).

**Figure 9.**
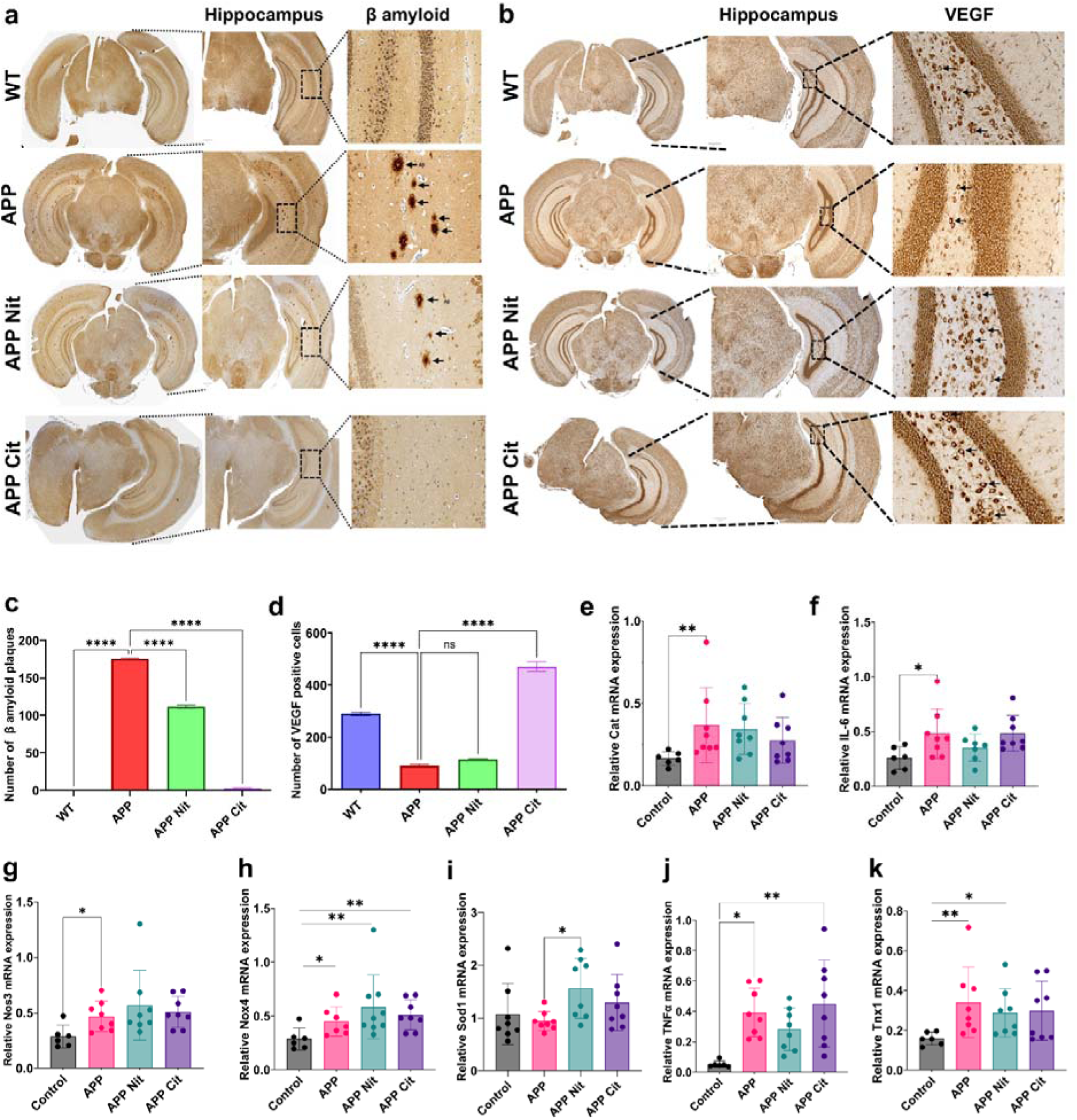
Hippocampal neurovascular pathology in APP/PS1 mice and adipose inflammatory/redox gene expression and the effects of citrulline and nitrite treatment. (a) Representative hippocampal IHC images for β-amyloid showing a significant increase in APP mice compared with WT and a significant reduction in both APP Cit and APP Nit, with APP Cit exhibiting the most significant decrease in β-amyloid burden. (b) Hippocampal VEGF IHC showing significantly reduced VEGF immunoreactivity in APP mice compared with WT and significantly increased VEGF staining in APP Cit and APP Nit relative to APP, with more substantial VEGF restoration in APP Cit. IHC quantification of (c) β-amyloid and (d) VEGF. RT-PCR analysis of gWAT showing inflammatory and redox gene expression for (e) Catalase (Cat), (f) Interleukin-6 (IL-6), (g) Nitric oxide synthase 3 (Nos3), (h) NADPH oxidase 4 (Nox4), (i) Superoxide dismutase-1 (Sod1), (j) Tumor necrosis alpha (TNF-α), and (k) thioredoxin 1 (Txn1).

Consistent with these peripheral molecular changes, immunohistochemical analysis of hippocampal brain sections demonstrated significant AD pathology and treatment-dependent effect (**Fig. 9a–d**). β-amyloid level was significantly increased in the hippocampus of APP mice compared with WT (**Fig. 9a, 9c**), while both APP Nit and APP Cit exhibited a significant reduction in β-amyloid burden relative to APP, with a more pronounced decrease observed in APP Cit. Conversely, VEGF expression was significantly reduced in the hippocampus of APP mice compared with WT (**Fig. 9b, 9d**) and was significantly restored in both APP Nit and APP Cit, again with a stronger effect in the citrulline-treated group.

## 4. Discussion

This study showed a coordinated disruption of adipose lipid metabolism, redox signaling, and mitochondrial function in APP/PS1 mice on high-fat diet (HFD). Our results revealed that nitric oxide-directed interventions with citrulline and nitrite partially normalize peripheral lipid–redox pathways and improve central neurovascular pathology. At the whole-body level, APP mice showed increased adiposity, glucose intolerance, reduced circulating glycerol, and increased adipocyte hypertrophy, consistent with impaired lipolysis and metabolic inflexibility reported in HFD-fed AD models [42, 43] . These systemic changes were consistent with the observed lipidomic profile, characterized by increased triglycerides and ceramide accumulation, accompanied by a broad reduction in major membrane phospholipids, including PC, PE, PG, LPC, and LPE. These changes suggest reduced phospholipid turnover and compromised mitochondrial membrane renewal, indicative of impaired membrane homeostasis [15, 44–46]. Neither citrulline nor nitrite significantly reduced obesity or adipocyte hypertrophy; however, citrulline showed a modest improvement in glucose handling at week 25, suggesting a partial restoration of adipose lipid remodeling [32, 47, 48].

In our study, untargeted and targeted lipidomics consistently indicated that APP adipose tissue exists in a pro-oxidative and pro-inflammatory microenvironment, characterized by increased levels of 13-HODE, 12-HETE, and 14-HDHA [49, 50]. Raman microscopy further demonstrated a shift from protective nitro-oleic acid toward excessive nitration of polyunsaturated fatty acids (nitro-CLA, nitro-AA, nitro-DHA), revealing a transition from regulated NO lipid signaling to pathological nitrative stress [31, 49, 50]. Citrulline and nitrite partially reversed these lipid changes, but through different mechanisms. Citrulline promoted restoration of nitro-oleic acid (nitro-OA) and supported PA → CDP-DAG → PC/PE phospholipid remodeling, whereas nitrite primarily acted through NOX4-linked redox signaling pathways [51, 52]. Consistently, RT-PCR analysis showed increased expression of TNF-α and NOX4, confirming a sustained inflammatory and oxidative state in APP adipose tissue [51].

In our study, FLIM analysis provided a functional metabolic correlation by demonstrating increased fluorescence lifetimes in APP, APP Cit, and APP Nit animals compared with WT, suggesting a shift toward increased protein-bound NADH and impaired mitochondrial redox cycling [20, 53]. These findings are consistent with the lipidomics data showing loss of mitochondrial supportive phospholipids, ceramide accumulation, and oxidized fatty acid production, which together impair electron transport chain organization and mitochondrial flexibility [54]. Although citrulline and nitrite improved the nitro-lipid balance and partially restored phospholipid metabolism, FLIM analysis showed that mitochondrial redox homeostasis was not fully recovered and remained lower than in WT.

Together, the RT-PCR and immunohistochemical findings suggest that redox imbalance and inflammatory activation in peripheral adipose tissue are closely associated with the development of neurovascular pathology in APP/PS1 mice. Increased expression of Nox4 across all APP groups indicates persistent activation of NADPH oxidase dependent redox signaling, consistent with lipidomic evidence of oxidative stress, ceramide accumulation, and nitro-lipid imbalance [54]. The relatively lower Nox4 induction in APP Cit compared with APP Nit suggests that citrulline may promote a more regulated redox environment than nitrite [55]. Increased TNF-α and IL6 expression, along with the upregulation of antioxidant genes such as Cat, Tnx1, and Sod1, indicate a chronic low-grade inflammatory state coupled with compensatory antioxidant responses, rather than an acute inflammatory activation, consistent with the unchanged Nos2 levels [51]. The selective upregulation of Sod1 in APP Nit mice suggests an increased oxidative load under nitrite treatment, consistent with the lipidomic profile showing enrichment of stress-associated lipid species in this group [32, 51].

Notably, these peripheral molecular changes were closely reflected in the hippocampus. APP mice displayed marked β-amyloid accumulation together with reduced VEGF expression, indicating impaired neurovascular support alongside enhanced amyloid pathology [56–58]. Treatment with either nitrite or citrulline significantly reduced amyloid burden and restored VEGF expression, indicating that targeting peripheral redox and inflammatory pathways can translate into measurable improvements in central neurovascular and amyloid pathology [59, 60]. The more pronounced reduction in β-amyloid and restoration of VEGF in APP Cit are consistent with lipidomic and nitro-lipid profiles indicating improved membrane remodelling, improved mitochondrial lipid composition, and lower nitrative stress with citrulline. Overall, these results support a model in which adipose redox–inflammatory signalling drives neurovascular dysfunction in APP/PS1 mice and suggest that citrulline is more effective than nitrite in restoring systemic redox balance and neurovascular integrity.

Our results showed an adipose lipid redox axis through which HFD worsens AD-related neurovascular injury. APP adipose tissue shows coordinated ceramide accumulation, oxidized lipid buildup, nitro-lipid imbalance, inflammatory activation, and disrupted mitochondrial redox metabolism. NO-targeted interventions partially reprogram these pathways, linking peripheral lipid redox remodelling to reduced hippocampal amyloid pathology and improved neurovascular signalling. Among them, citrulline more effectively restores phospholipid and nitro-lipid homeostasis, highlighting its potential to target peripheral metabolic inflammation in neurodegeneration.

## 5. Conclusions

This study identifies adipose tissue as a key interface linking metabolic stress, redox imbalance, and neurovascular pathology in Alzheimer’s disease. In HFD-fed APP/PS1 mice, adipose lipid remodelling was changed, with reduced phospholipid turnover, ceramide and storage lipid accumulation, disrupted oxidized lipid mediator profiles, and impaired nitro-fatty acid metabolism. These peripheral disturbances coincided with sustained oxidative and inflammatory signalling, mitochondrial redox dysfunction, increased hippocampal amyloid deposition, and loss of VEGF-mediated neurovascular support. Integrative analyses combining lipidomics, Raman imaging, FLIM, and molecular and histological assessments uncovered an adipose–brain lipid redox axis in AD. NO-based interventions partially reprogrammed this axis, with citrulline more effectively restoring membrane lipid remodeling and nitro-lipid signaling, whereas nitrite predominantly influenced stress-associated lipid pathways. Overall, these findings underscore the role of peripheral lipid–redox homeostasis in modulating neurovascular resilience and suggest that targeting adipose NO–lipid interactions may help reduce metabolic contributions to neurodegeneration.

## Acknowledgment

We thank Dr. Fabrizio Donnarumma for the help with the untargeted lipidomics experiments. We thank Dr. David Burk for the help with the FLIM experiments. We thank Dr. Natalia M. Stephanes for the help with plotting the data.

## Funding agency

Research reported in this publication was supported by the National Institute of General Medical Sciences of the National Institutes of Health under award number R35GM150564. W.N.B. acknowledges the support of R35GM154857.

