## Supporting Information for "Integrated Lipidomics and Nitro-Fatty Acid Profiling Link Adipose Redox Imbalance to Alzheimer’s Disease-Related Neurovascular Injury"

**Table S3.** Primers used for the RT-PCR in the study

| **Primer** | **Species** | **Forward** | **Reverse** |
| --- | --- | --- | --- |
| Nos-3 | Mouse | CTTGAGGATGTGGCTGTGT | TGGTCCACTATGGTCACTTTG |
| Sod-1 | Mouse | GGTTCCACGTCCATCAGTATG | GTCCTTTCCAGCAGTCACAT |
| Txn1 | Mouse | GGTGTGGACCTTGCAAAATGATC | GGCTTCAAGCTTTTCCTT |
| Tnfa | Mouse | AAGCCTGTAGCCCACGTCGTA | AGGTACAACCCATCGGCTGG |
| Nox-4 | Mouse | TCCTACTGAAACCAAAGCAACA | AATGAAGGGCAGAATCTCAGAG |
| Cat | Mouse | CAAGTTGGTTAATGCAGATGGAG | ATCTTCCTGAGCAAGCCTTC |
| IL-6 | Mouse | TCCATCCAGTTGCCTTCTTG | TTCCACGATTTCCCAGAGAAC |

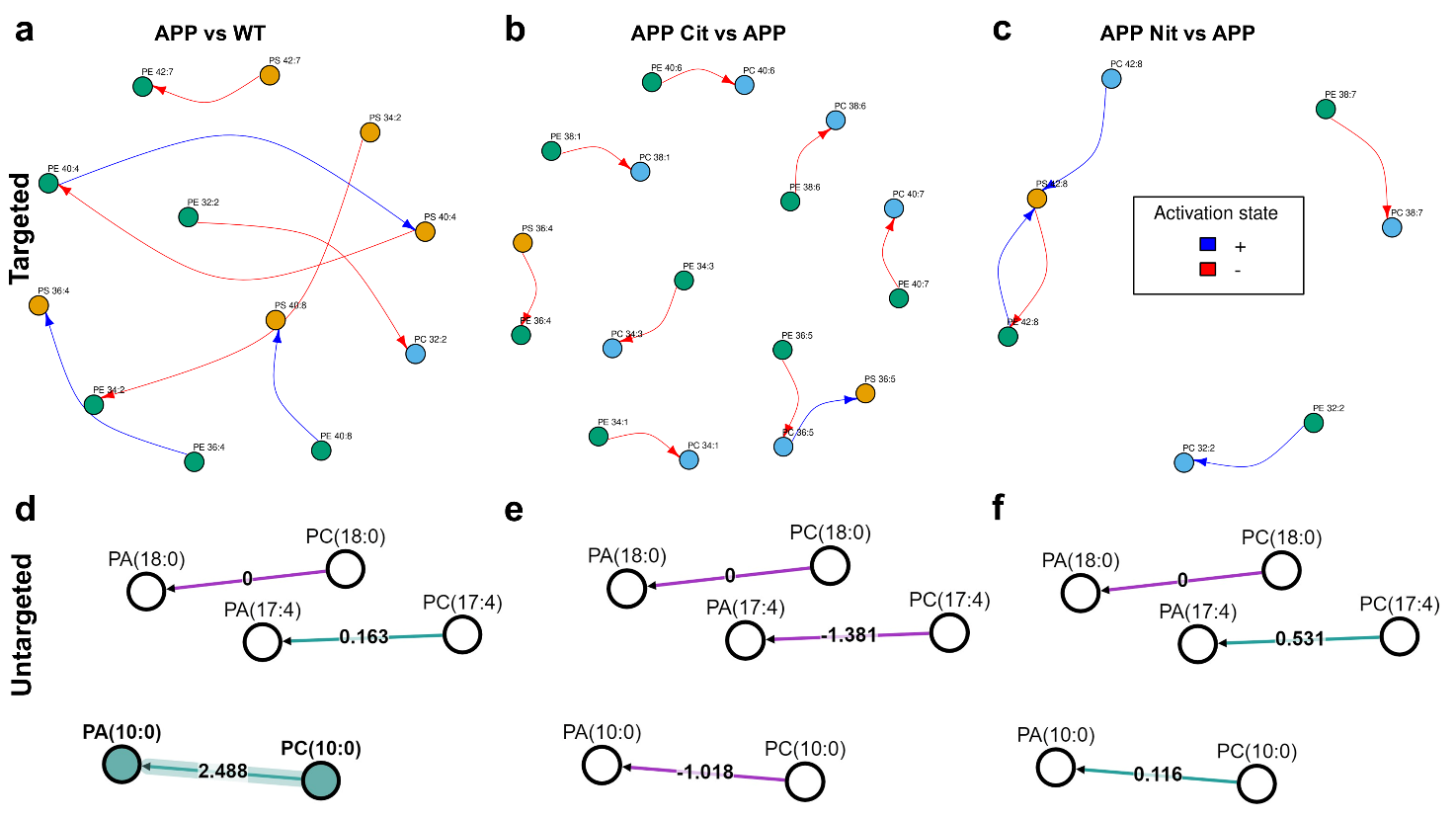

**Figure S1.** (a–c) LipidOne‑based pathway maps highlight activation of Ptdss1/2‑mediated PE→PS steps and suppression of Psd and Pemt in APP vs WT and APP Cit vs APP, whereas APP Nit vs APP further promotes formation of PS 42:8 while limiting its reconversion to PE or PC. (d–f) BioPAN analysis of PA→PC pairs shows PA(10:0)→PC(10:0) and PA(17:4)→PC(17:4) are strongly upregulated in APP vs WT, partly preserved in APP vs APP Nit, and suppressed in APP vs APP Cit, indicating specific PA–PC conversions that are APP‑driven and preferentially reduced by citrulline.

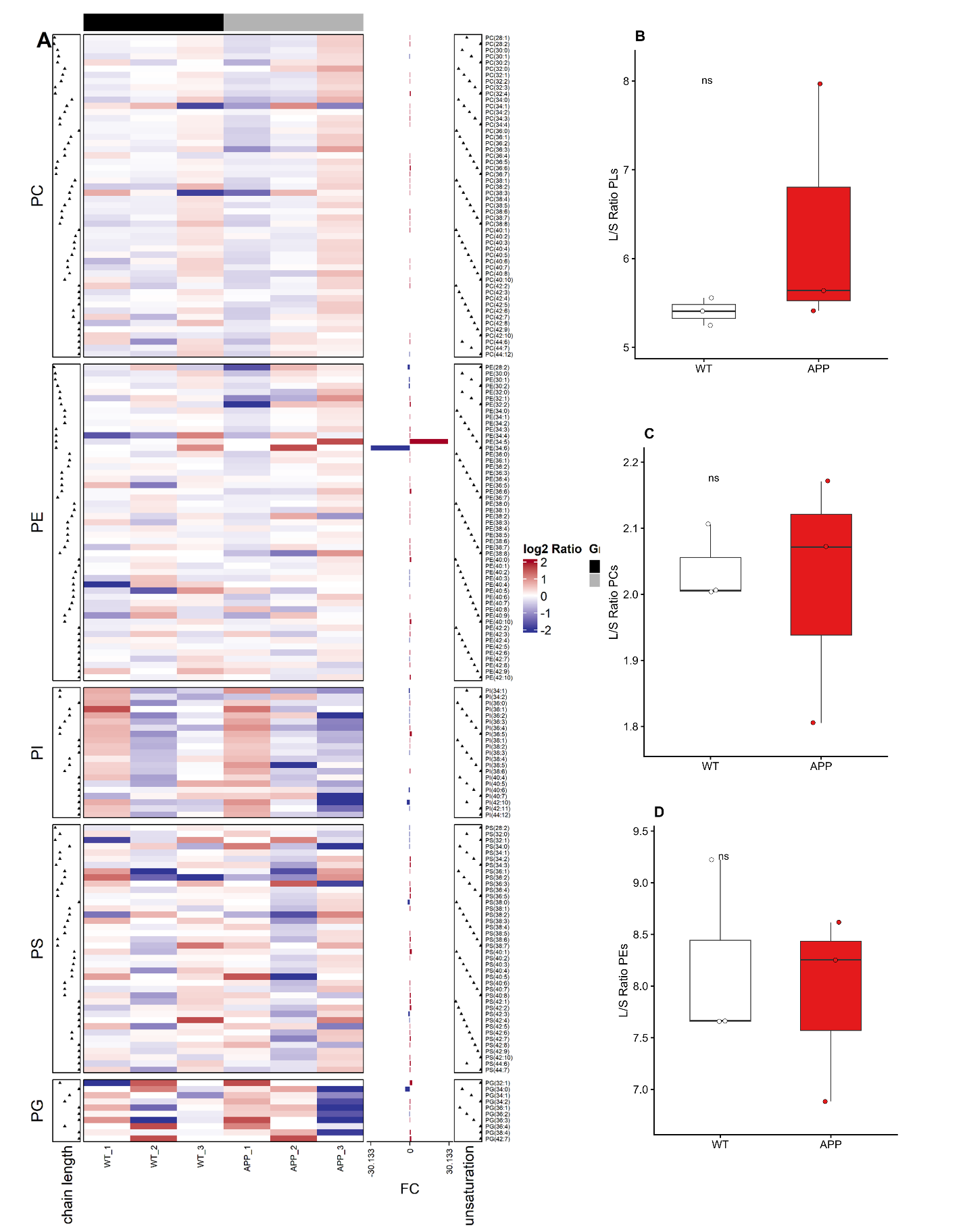

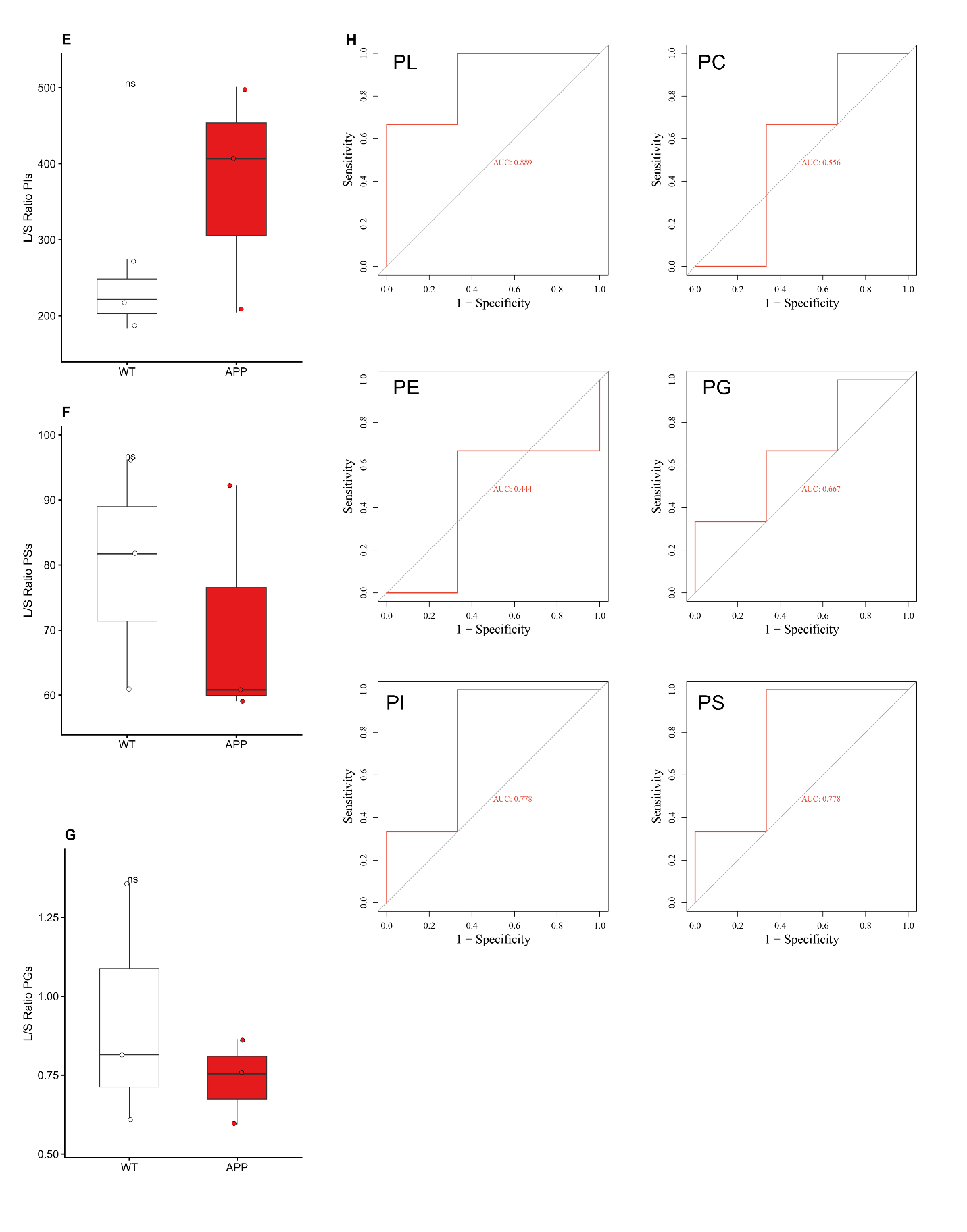

**Figure S2.** Comparative lipidomic profiling and discriminatory potential of phospholipid species in APP transgenic mice. (A) Heatmap and structural distribution of phospholipids. Differential abundance of individual lipid species across Phosphatidylcholine (PC), Phosphatidylethanolamine (PE), Phosphatidylinositol (PI), Phosphatidylserine (PS), and Phosphatidylglycerol (PG) classes in Wild-Type (WT, n=3) and APP transgenic (APP, n=3) mice. Color scale represents the log2 ratio of abundance. Side panels indicate the structural characteristics of each species: chain length (left) and degree of unsaturation (right). The center bar plot (FC) indicates the fold-change for specific highly differential species. (B–G) Long-to-short (L/S) chain ratio analysis. Box plots represent the ratio of long-chain to short-chain fatty acids within specific phospholipid classes: (B) Total Phospholipids (PLs), (C) PCs, (D) PEs, (E) PIs, (F) PSs, and (G) PGs. While the APP group exhibits a trend toward increased L/S ratios in total PLs and PIs, differences did not reach statistical significance (ns, p>0.05 via Student’s t-test). Data points represent individual biological replicates; horizontal lines indicate the median. (H) Receiver Operating Characteristic (ROC) analysis. ROC curves evaluating the performance of L/S ratios as biomarkers to discriminate between WT and APP genotypes. Area Under the Curve (AUC) values are provided for total PLs ($\text{AUC}=0.889$AUC=0.889), PCs (AUC=0.556), PEs (AUC=0.444), PGs (AUC=0.667), PIs (AUC=0.778), and PSs (AUC=0.778). The high AUC for total PLs and PIs suggests these specific lipid metrics have the highest diagnostic potential in this Alzheimer’s disease model.

**.**

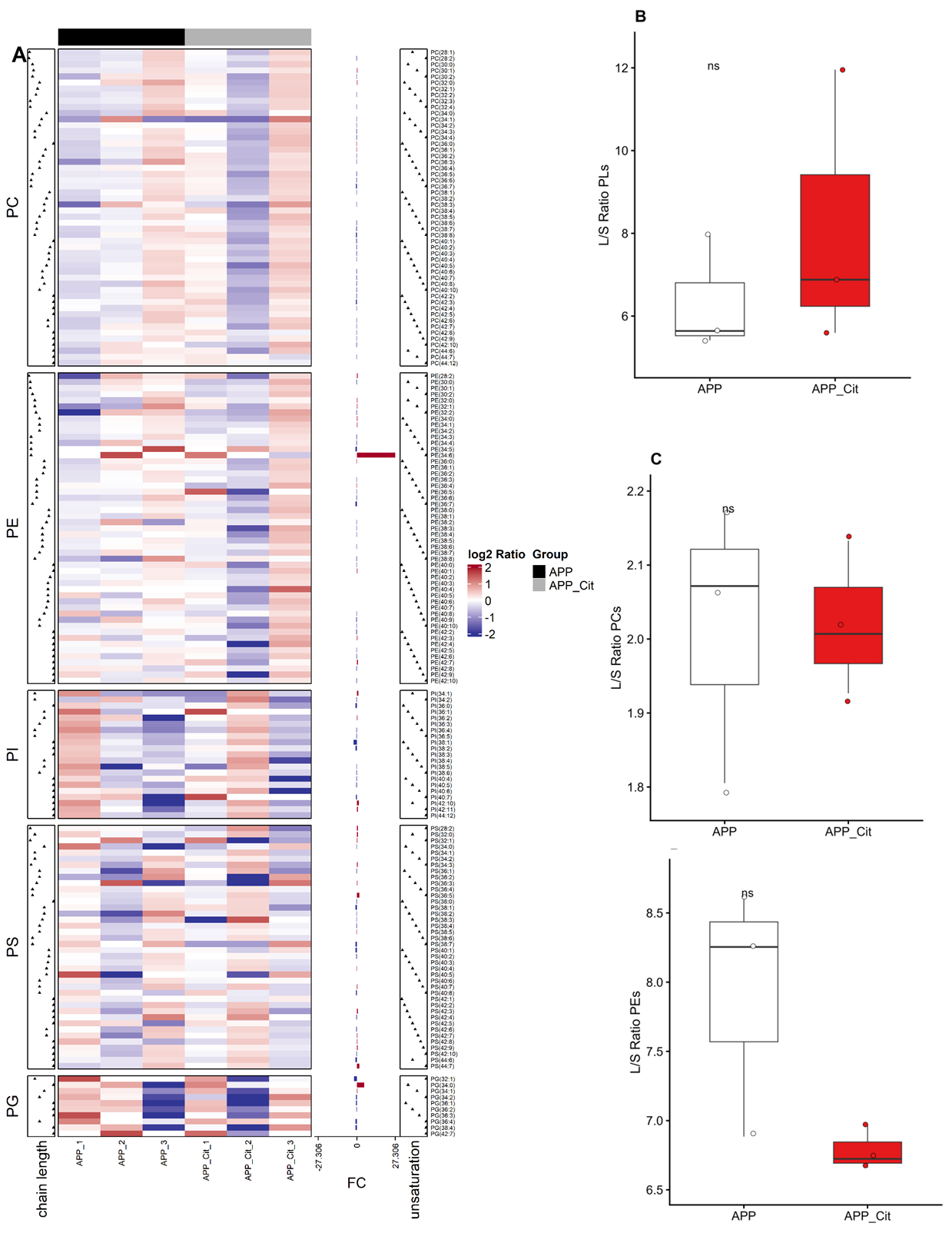

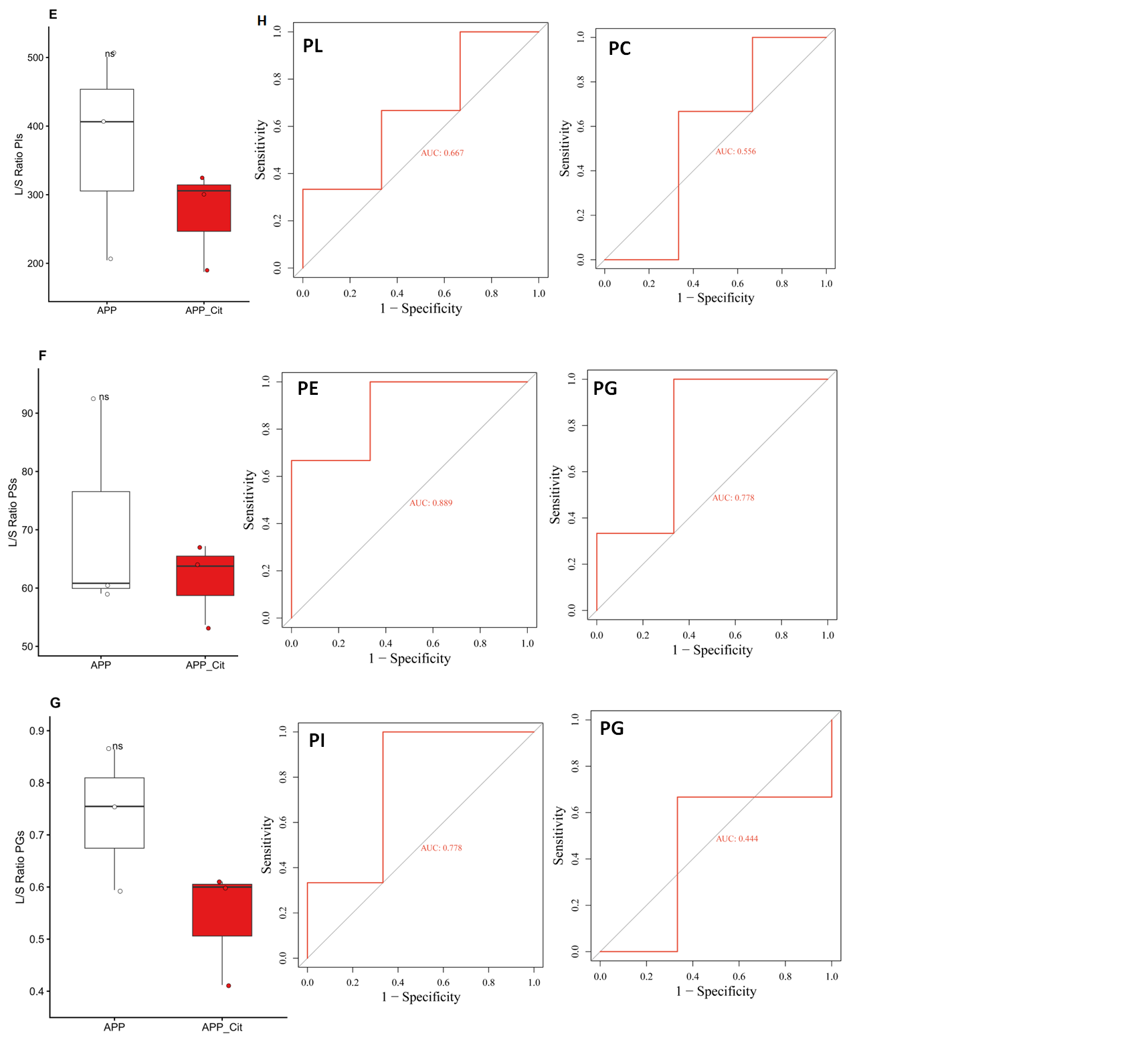

**Figure S3**. Lipidomic remodeling and diagnostic performance of phospholipid chain-length ratios in Citrate-treated APP mice. (A) Heatmap of phospholipid species distribution. Heatmap illustrating the relative abundance of individual lipid species within the Phosphatidylcholine (PC), Phosphatidylethanolamine (PE), Phosphatidylinositol (PI), Phosphatidylserine (PS), and Phosphatidylglycerol (PG) classes. Comparisons are shown between untreated APP transgenic mice (APP, n=3, black bar) and Citrate-treated APP mice (APP_Cit, n=3, grey bar). The color gradient represents the log2 ratio of abundance. Accompanying dot plots indicate the chain length and degree of unsaturation for each species. The central bar plot (FC) highlights the fold-change of specific discriminatory metabolites. (B–G) Longitudinal chain-length (L/S) ratio analysis. Box-and-whisker plots comparing the ratio of long-chain to short-chain fatty acids (L/S Ratio) between APP (white) and APP_Cit (red) groups across various classes: (B) Total Phospholipids (PLs), (C) PCs, (D) PEs, (E) PIs, (F) PSs, and (G) PGs. Citrate treatment shows a trend toward reducing the L/S ratio in PE, PI, and PG classes, although these changes did not reach statistical significance (ns, p>0.05 via Student’s t-test). Data points represent individual biological replicates; horizontal lines indicate the median. (H) ROC curve analysis for treatment discrimination. Receiver Operating Characteristic (ROC) curves evaluating the sensitivity and specificity of L/S ratios in discriminating the Citrate-treated group from the untreated APP group. Area Under the Curve (AUC) values are indicated for total PLs (AUC=0.667), PCs (AUC=0.556), PEs (AUC=0.889), PGs (AUC=0.778 and 0.444 for different subsets), PIs (AUC=0.778), and PSs (AUC=0.778). The high AUC for PE suggests that the long-to-short chain ratio in this lipid class is the most robust metabolic indicator of the Citrate treatment effect.

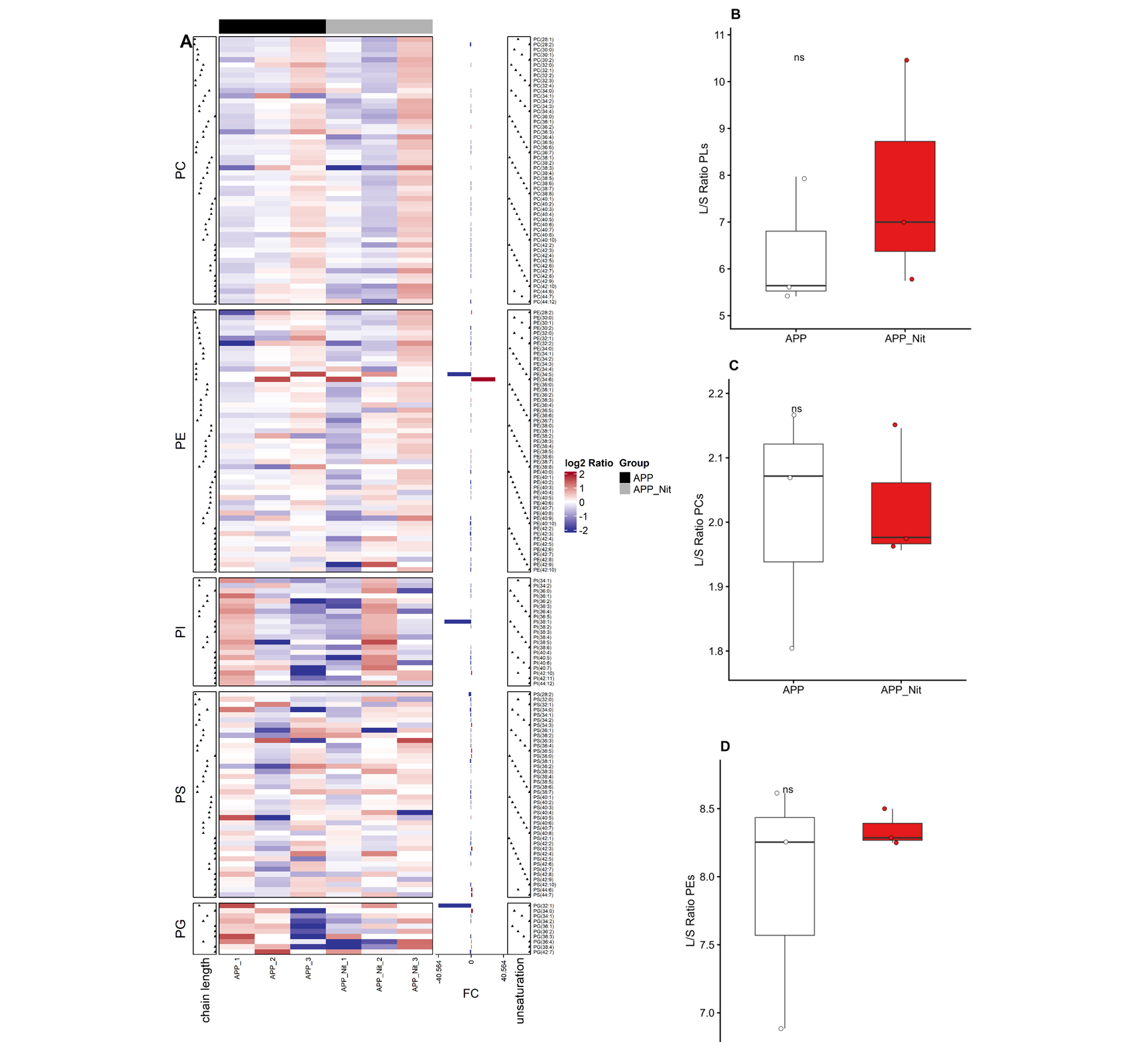

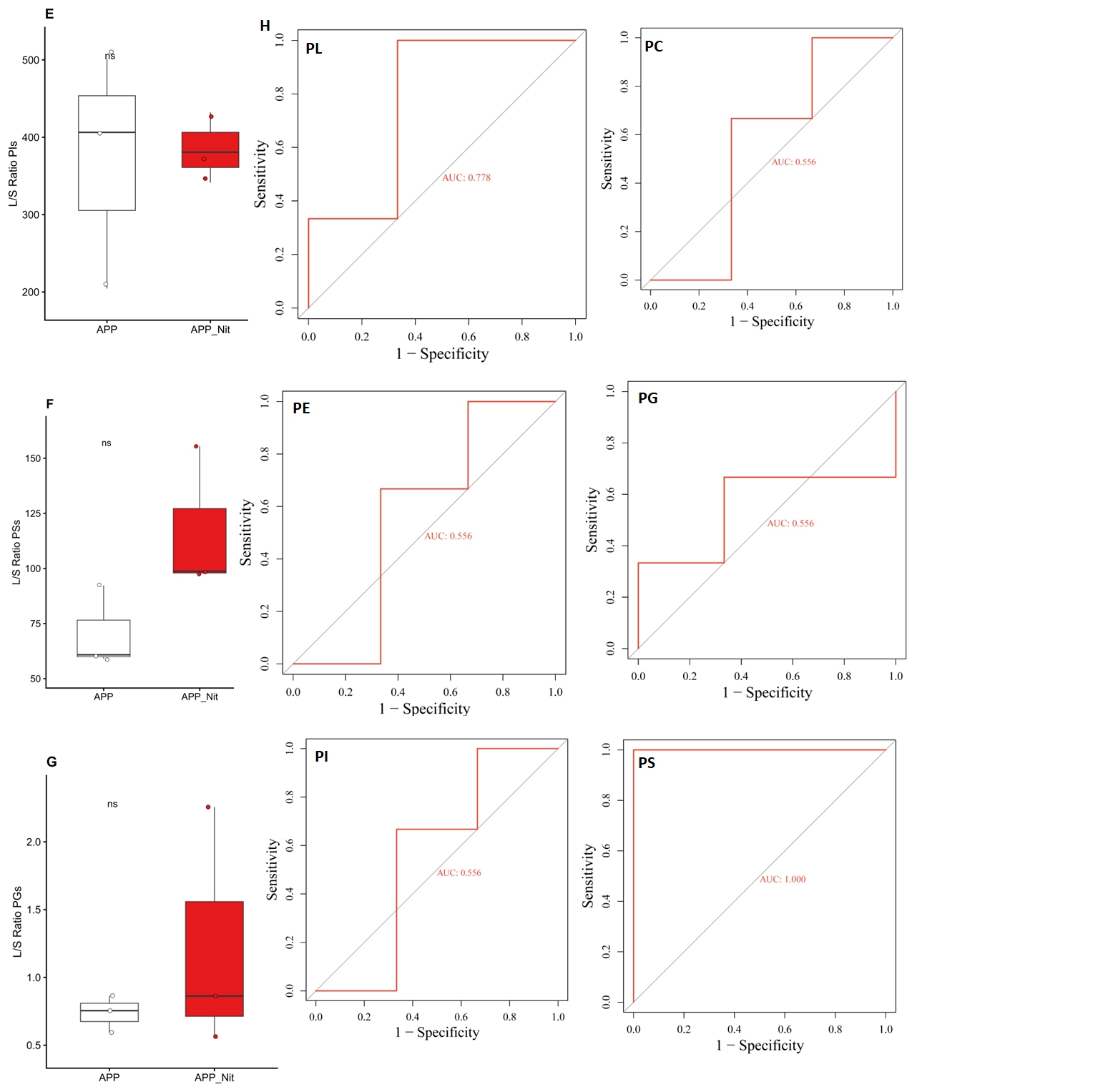

**Figure S4.** Lipidomic remodeling and classification performance of phospholipid chain-length ratios in Nitrate-treated APP transgenic mice. (A) Heatmap of phospholipid species distribution and structural characteristics. Relative abundance of individual lipid species across Phosphatidylcholine (PC), Phosphatidylethanolamine (PE), Phosphatidylinositol (PI), Phosphatidylserine (PS), and Phosphatidylglycerol (PG) classes. The heatmap compares untreated APP transgenic mice (APP, n=3, black bar) with Nitrate-treated APP mice (APP_Nit, n=3, grey bar). Color intensity represents the $\log_{2}$log2 ratio of abundance. Marginal dot plots indicate the chain length (left) and unsaturation degree (right) for each species. The central bar plot denotes the fold-change (FC) for specific discriminatory lipid species. (B–G) Long-to-short (L/S) chain ratio analysis across lipid classes. Box-and-whisker plots illustrating the ratio of long-chain to short-chain fatty acids in the APP (white) and APP_Nit (red) groups for: (B) Total Phospholipids (PLs), (C) PCs, (D) PEs, (E) PIs, (F) PSs, and (G) PGs. Nitrate treatment resulted in a trend toward increased L/S ratios in total PLs and PSs, though these differences were not statistically significant (ns, p>0.05 via Student’s t-test). Data points represent individual biological replicates; horizontal lines indicate the median. (H) ROC curve analysis for treatment discrimination. Receiver Operating Characteristic (ROC) curves evaluating the diagnostic potential of L/S ratios to distinguish between Nitrate-treated and untreated APP mice. Area Under the Curve (AUC) values are provided for total PLs (AUC=0.778), PCs (AUC=0.556), PEs (AUC=0.556), PGs (AUC=0.556), PIs (AUC=0.556), and PSs (AUC=1.000). The perfect AUC score for PS suggests that the long-to-short chain ratio in this specific phospholipid class is a highly robust biomarker for the metabolic effects of Nitrate treatment in this model.

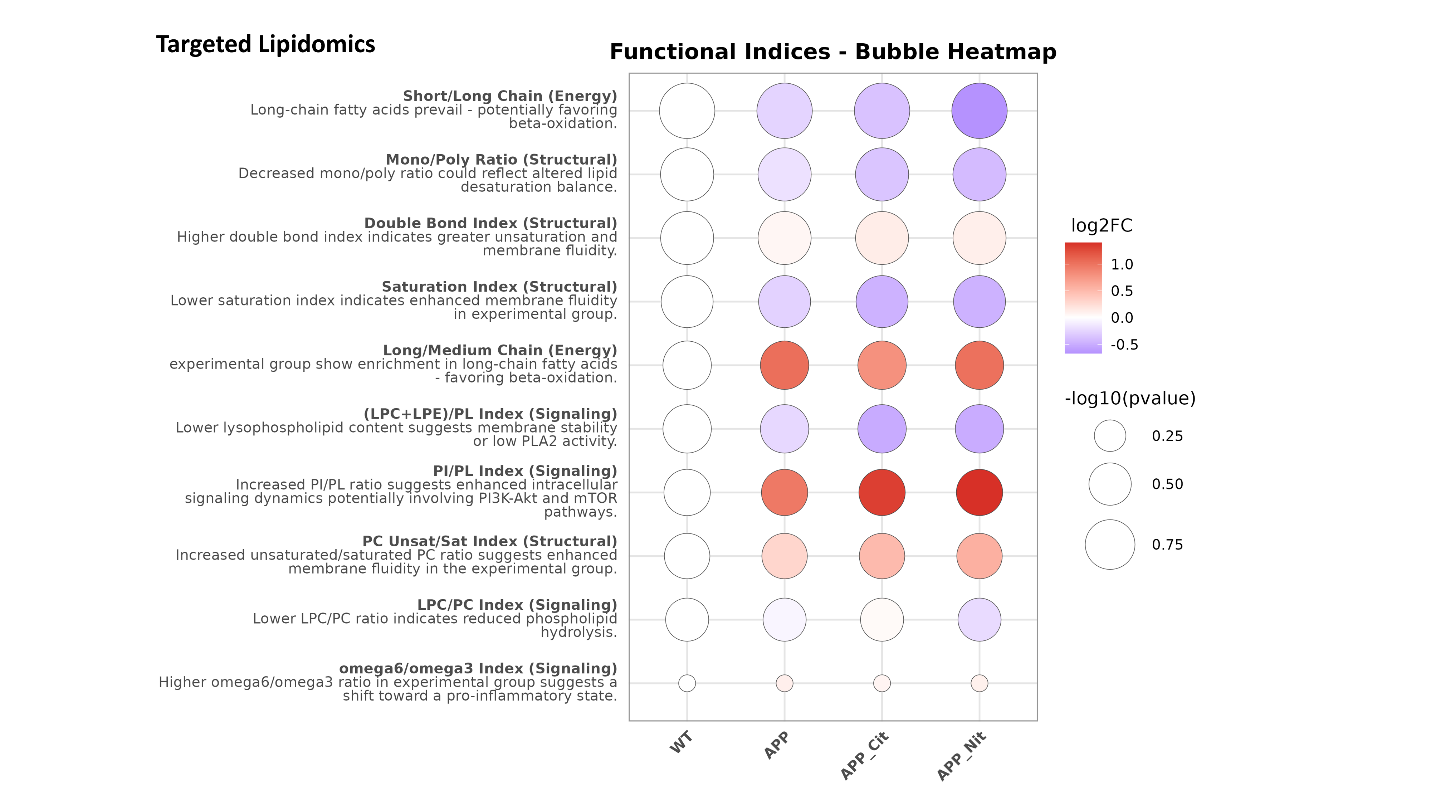

**Figure S5.** Targeted lipidomics and functional indices of membrane homeostasis in APP transgenic mice under Nitrate and Citrate treatment. Bubble heatmap illustrating the metabolic and structural shifts in the brain lipidome across Wild-Type (WT), APP transgenic (APP), Citrate-treated APP (APP_Cit), and Nitrate-treated APP (APP_Nit) groups. Functional indices are categorized into Energy, Structural, and Signaling domains to provide a holistic view of lipid remodeling.

Functional Indices and Biological Implications:

- Energy Metabolism: Shifts in the Short/Long Chain and Long/Medium Chain indices suggest a prevalence of long-chain fatty acids in the APP and treated groups, potentially favoring mitochondrial β-oxidation pathways.
- Structural Integrity and Fluidity: The Double Bond Index and PC Unsat/Sat Index indicate a trend toward increased unsaturation in experimental groups, while the Saturation Index and Mono/Poly Ratio reflect alterations in lipid desaturation balance, collectively suggesting modifications in membrane fluidity.
- Signaling Dynamics: The significant increase in the PI/PL Index across APP and treated groups suggests enhanced intracellular signaling, potentially involving the PI3K-Akt and mTOR pathways. Conversely, the reduction in the (LPC+LPE)/PL and LPC/PC indices in the treated groups suggests improved membrane stability or reduced Phospholipase A2 (PLA2) activity and phospholipid hydrolysis.
- Inflammatory State: The ω-6/ω-3 ratio serves as a proxy for the pro-inflammatory environment, with higher ratios indicating a shift toward neuroinflammatory signaling.

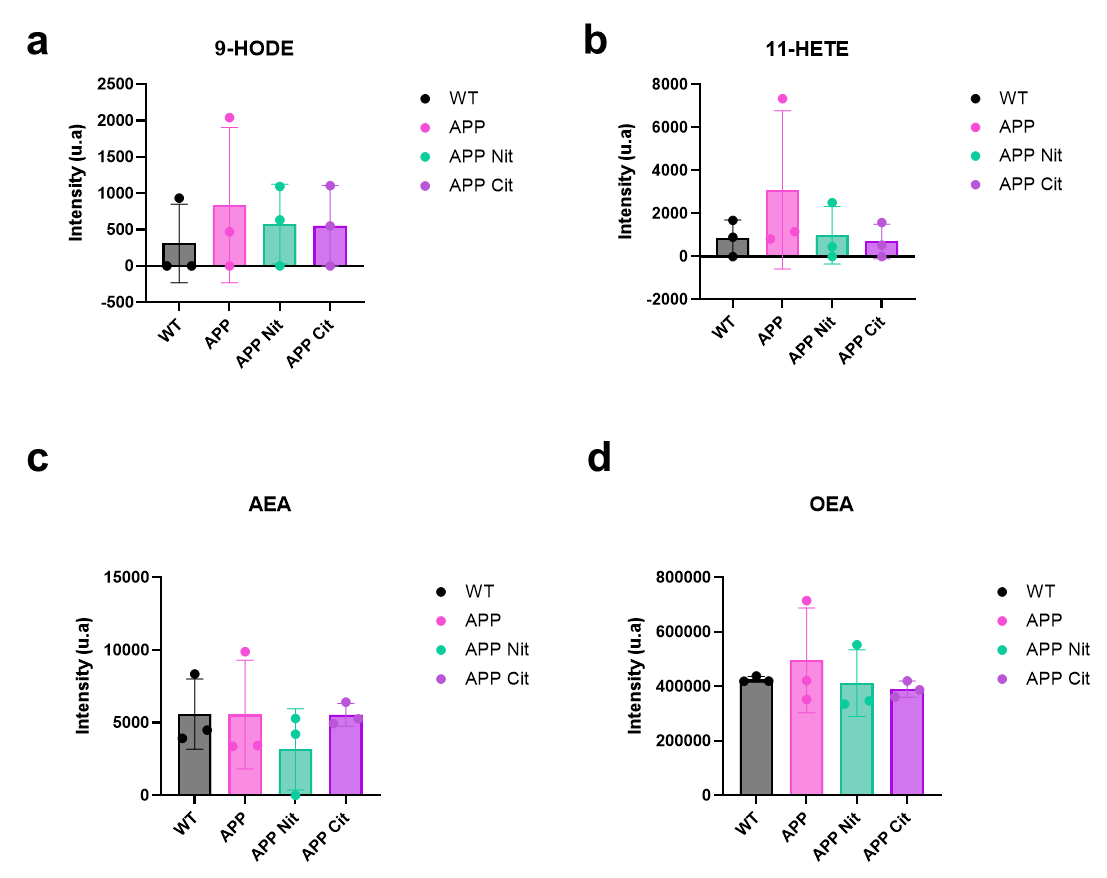

**Figure S6.** Lipidomic analysis of HODEs, HETEs, and endocannabinoid-related ethanolamides in APP transgenic mice. Quantification of specific lipid species in brain tissue (or specify sample type, e.g., cortical homogenates) across four experimental groups: Wild-Type (WT), APP transgenic (APP), and APP transgenic mice treated with Nitrate (APP Nit) or Citrate (APP Cit). (a) Levels of 9-HODE (9-hydroxyoctadecadienoic acid), a marker of lipid peroxidation and oxidative stress. (b) Levels of 11-HETE (11-hydroxyeicosatetraenoic acid), a non-enzymatic oxidation product of arachidonic acid. (c) Levels of AEA (Anandamide/N-arachidonoylethanolamine), a primary endocannabinoid neurotransmitter. (d) Levels of OEA (Oleoylethanolamide), a lipid mediator involved in metabolic regulation and neuroprotection. Data are presented as individual biological replicates ($n=3\text{–}4$ per group) with bars representing the mean $\pm$ SD. Intensity is expressed in arbitrary units (a.u). Statistical analysis (e.g., one-way ANOVA with Tukey’s post-hoc test) revealed no significant differences ($p>0.05$) between groups for these specific metabolites, despite a trend toward increased variance in the untreated APP group.

**Table S4.** Raman peak assignments

| **Species** | **Diagnostic Raman peaks (cm⁻¹)** | **Raman assignment** | **Notes** |
| --- | --- | --- | --- |
| **9-Nitrooleate (9-NO₂-OA)** | **1339** | **ν_s_(NO₂)** (nitro symmetric stretch) | Clear nitro marker; no aromatic ring peaks. Can overlap with CH wag/twist region but usually stands out. |
|  | 1440 | CH₂ scissoring | Strong lipid backbone marker (useful for normalization of spectra). |
|  | **1525** | **ν_as_(NO₂)** (nitro asymmetric stretch) | Second nitro marker (weak). |
|  | 1667 | ν(C=C) (alkene stretch; nitroalkene tends to shift upward vs native OA) | Typically, one main C=C feature, moderate width are present. |
|  | 2856, 2935 | CH₂/CH₃ stretches | Strong lipid region. |
|  | 3010–3030 | =C–H stretch | Present but weaker than polyunsaturated chains. |
| **9(E),11(E)-9-nitro-CLA** | **1319** | **ν_s_(NO₂)** | Present (lower frequency compared to NO₂-OA). |
|  | 1445; 2856; 2922 | CH₂ scissor; CH stretches | Lipid |
|  | **1515** | **ν_as_(NO₂)** | Present (but weak). |
|  | **1649** | **ν(C=C)** of **conjugated diene / nitroalkene** | **Best discriminator** vs **9-NO₂-OA**: (strong). |
|  | 3011 | =C–H stretch | Weak |
| **Arachidonoyl p-nitroaniline (nitro-AA)** | **1014** | **Aromatic ring breathing** | Sharp, strong |
|  | **1599** | Aromatic ring C=C stretch | Sharp aromatic band(s), strong. |
|  | **1341** | ν_s_(NO₂) on aromatic ring | Nitro present; very strong. |
|  | **1510; 1550** | ν_as_(NO₂) (overlaps with Amide II region) | Broadened/complex feature because of overlap. |
|  | **1658** | **Amide I** (C=O stretch) ± lipid ν(C=C) | **Amide I** and lipid C=C region overlap → broadened/shifted peak. |
|  | 1550 | **Amide II** (N–H bend/C–N stretch) | Merge with νas(NO₂). |
|  | 1257 | **Amide III** | Protein |
|  | 2902; 3017 | CH stretches; =C–H | Lipid chain related. |
| **p-Nitrophenyl sulfonamide DHA (nitro-DHA)** | **1110; 1175** | **ν_s_(SO₂)** (sulfonyl symmetric stretch) | Weak but distinct |
|  | **1352** | **ν_as_(SO₂)** + ν_s_(NO₂) overlap region | Very strong; |
|  | **1020** | Aromatic ring breathing | Weak aromatic (nitrophenyl) mode. |
|  | **1591** | Aromatic ring C=C stretch | Sharp. |
|  | **1535** | ν_as_(NO₂) | Nitro (aromatic) mode. |
|  | 1658 (broad) | ν(C=C) from DHA (multiple double bonds) | PUFA gives broader C=C region; still may be influenced by nearby functional group environment. |
|  | 3017 (stronger) | =C–H stretch | DHA typically shows stronger olefinic CH than monoenes. |
|  | 2873; 2913; 1443 | CH stretches; CH₂ scissor | Strong lipid backbone. |
